# The histone variant H2A.Z and chromatin remodeler BRAHMA act coordinately and antagonistically to regulate transcription and nucleosome dynamics in *Arabidopsis*

**DOI:** 10.1101/243790

**Authors:** E. Shannon Torres, Roger B. Deal

**Author notes:** Correspondence: Roger B. Deal.

## Abstract

Plants adapt to changes in their environment by regulating transcription and chromatin organization. The histone H2A variant H2A.Z and the SWI2/SNF2 ATPase BRAHMA have overlapping roles in positively and negatively regulating environmentally responsive genes in *Arabidopsis*, but the extent of this overlap was uncharacterized. Both have been associated with various changes in nucleosome positioning and stability in different contexts, but their specific roles in transcriptional regulation and chromatin organization need further characterization. We show that H2A.Z and BRM act both cooperatively and antagonistically to contribute directly to transcriptional repression and activation of genes involved in development and response to environmental stimuli. We identified 8 classes of genes that show distinct relationships between H2A.Z and BRM and their roles in transcription. We found that H2A.Z contributes to a range of different nucleosome properties, while BRM stabilizes nucleosomes where it binds and destabilizes and/or repositions flanking nucleosomes. H2A.Z and BRM contribute to +1 nucleosome destabilization, especially where they coordinately regulate transcription. We also found that at genes regulated by both BRM and H2A.Z, both factors overlap with the binding sites of light-regulated transcription factors PIF4, PIF5, and FRS9, and that some of the FRS9 binding sites are dependent on H2A.Z and BRM for accessibility. Collectively, we comprehensively characterized the antagonistic and cooperative contributions of H2A.Z and BRM to transcriptional regulation, and illuminated their interrelated roles in chromatin organization. The variability observed in their individual functions implies that both BRM and H2A.Z have more context-specific roles within diverse chromatin environments than previously assumed.

## INTRODUCTION

As sessile organisms, plants have evolved a plethora of physiological responses to endure adverse environmental conditions. External signals are often transmitted to the nucleus, triggering a transcriptional response network that facilitates a multidimensional response to the external stimuli (Rymen and Sugimoto 2012; Barah et al., 2016, Urano et al., 2010) In eukaryotes, DNA associates with histones and other nuclear proteins to form a highly condensed chromatin structure. The arrangement of these proteins can either facilitate or obstruct transcription factor (TF) binding to target regulatory sequences, and therefore impact the ability of transcriptional machinery to modulate these transcriptional responses (Weber et al., 2014; Lai and Pugh 2017).

At its most basic level, chromatin is composed of DNA wrapped around an octamer of histone proteins to form nucleosomes (Luger et al., 1997). Chromatin-binding proteins such as histone post-translational modifying enzymes and chromatin remodelers interact with nucleosomes and influence their positioning, stability, and ability to interact with other proteins, thus regulating DNA accessibility (Vergara and Gutierrez 2017). Chromatin remodeling complexes (CRCs) use the energy of ATP to disrupt the interactions between DNA and histones in order to evict nucleosomes, eject histone dimers, slide nucleosomes, or exchange canonical histones for variant forms (Narlikar et al., 2013; Clapier et al., 2017). Chromatin remodeling is a key part of regulating genome stability, DNA replication, DNA damage repair, and transcription, which in turn affects development, homeostasis, and how an organism responds to changes in the environment (Probst and Mittelsten Scheid 2015; Vergara and Gutierrez 2017).

The combined effects of many chromatin-regulating proteins at a locus create opposing and redundant forces that maintain proper transcription level and integrate a myriad of endogenous and exogenous signals (Vergara and Gutierrez 2017). One way to define the individual contributions of chromatin regulating factors *in vivo* is to evaluate how such proteins coordinately or antagonistically contribute to chromatin organization and transcription. Studies in *Arabidopsis thaliana* evaluating the histone H2A variant H2A.Z, which is deposited by the SWR1 CRC, and the SWI2/SNF2 CRC separately have revealed that they both modulate chromatin organization and transcription to regulate developmental processes and responses to the environment (Wu et al., 2015; Lee and Seo 2017). More directly, Farrona et al. (2011) proposed that the SWI2/SNF2 ATPase BRAHMA (BRM) and the incorporation of H2A.Z into chromatin by the SWR1 CRC have antagonistic roles to modulate *Flowering Locus C (FLC)* transcription levels and the developmental timing of flowering. In yeast, mutations in H2A.Z increase dependence on the SWI2/SNF2 complex for transcriptional activation of several genes, implying that the histone variant and chromatin remodeler have cooperative functions (Santisteban et al., 2000). Furthermore, in mammals, SWI2/SNF2 subunits interact with H2A.Z, although the implications of this interaction have not been explored (Li et al., 2012; Zhang et al., 2017b). While the SWR1 CRC and the SWI2/SNF2 complex have parallel roles in development and environmental responses in plants, there is a dearth of studies that focus on the direct intersection of these two complexes in chromatin and transcriptional regulation. Since an antagonistic relationship was already established between H2A.Z and BRM at the *Arabidopsis FLC* gene, we decided to characterize the extent to which these two factors interact in chromatin organization and transcriptional regulation.

In this study, we demonstrate that H2A.Z and BRM interact to impact nucleosome organization and regulate transcription across the *Arabidopsis* genome. We assessed the overall and direct transcriptional contributions of H2A.Z and BRM by performing transcriptional profiling in combination with BRM and H2A.Z localization information in wild type, single, and double mutants. Using mutants lacking BRM, mutants for the ARP6 subunit of the SWR1 CRC that are defective in H2A.Z incorporation into chromatin, or double mutants depleted for both, we identified 8 different classes of co-targeted genes where transcription is coordinately or antagonistically regulated by H2A.Z and BRM, including genes that are up- or down-egulated only in double mutants. The genes regulated by both H2A.Z and BRM contribute to a number of biological processes, including development and responses to various stimuli. By experimentally verifying that these genes are direct targets of H2A.Z or BRM, the regulatory relationships we identified allude to cooperative and antagonistic functions between BRM and H2A.Z in chromatin regulation and transcription.

We further explored how BRM and H2A.Z contribute to nucleosome organization to facilitate these transcriptional changes by measuring nucleosome occupancy and positioning. We found that BRM is involved in nucleosome stabilization at nucleosome-depleted regions (NDR), both distal and proximal to the transcription start sites (TSS), and contributes to destabilization and/or repositioning of flanking nucleosomes. On the other hand, H2A.Z-containing nucleosomes show highly variable changes in nucleosome properties upon H2A.Z depletion. At loci where both H2A.Z and BRM are found together in the genome, BRM usually destabilizes nucleosomes, especially the +1 nucleosome, while H2A.Z can also destabilize +1 nucleosomes at some loci. In addition, we identified binding sites of light-responsive TFs PIF4, PIF5, and FRS9 that are enriched at BRM and H2A.Z co-targeted genes, and show that nucleosome occupancy is dependent on BRM and H2A.Z at some FRS9 binding sites. These findings point to a role for both H2A.Z and BRM in regulating nucleosome positioning and stability in coordinately and antagonistically regulated genes involved in light response and other responses to stimuli. Collectively, our findings indicate that the relationship between BRM and H2A.Z is more complex than solely antagonizing or complementing the chromatin organizing function of the other, and our datasets will be useful for future studies to explore the contexts in which BRM and H2A.Z contribute to chromatin organization or transcriptional regulation.

## RESULTS

### Analysis of transcriptional changes in *arp6* and *brm* mutants

Both H2A.Z and BRM contribute to transcriptional repression and transcriptional activation, but it is unknown how the presence of one might affect the role of the other in transcriptional regulation. (Marques et al., 2010; Archacki et al., 2016). To identify the genes in *Arabidopsis* that are transcriptionally regulated by BRM and H2A.Z, we performed RNA sequencing (RNA-seq) experiments. We used plants with the *brm-1* allele, since it was previously characterized as a BRM null mutant (Hurtado et al., 2006). To study plants with H2A.Z-depleted nucleosomes, we used null mutants for the ARP6 component of the SWR1 chromatin remodeling complex (CRC) which is primarily responsible for incorporating H2A.Z into nucleosomes (Deal et al., 2007). In *Arabidopsis*, three genes encode the pool of H2A.Z proteins and there are no completely null triple H2A.Z mutants available, which complicates genetic work (Coleman-Derr and Zilberman 2012). Other studies have verified that ARP6 is required for proper H2A.Z incorporation into nucleosomes and *arp6* mutants phenocopy H2A.Z mutants, making *arp6* mutants a logical proxy for H2A.Z mutants in our genetic study (Sura et al., 2017; March-Diaz et al., 2008; Berriri et al., 2016). Therefore, we identified the genes that are differentially expressed (DE) in *brm-1* mutants, *arp6–1* mutants, and *arp6–1;brm-1* double mutants compared to wild type (WT) plants. We focused our analyses on genes that had >1.5x fold change at a false discovery rate of <0.2. We chose this less stringent cutoff so as to avoid excluding true positives while describing general processes regulated by these factors. We later use additional criteria to identify specific genes for downstream analyses. RNA was isolated from above soil, green tissue from developmentally staged plants with 4–5 leaves. We collected tissue based on developmental stage instead of age of the plant because *brm* and *arp6;brm* double mutants present delayed developmental progression (Boyes et al., 2001;Hurtado et al., 2006). After performing RNA-seq, we identified 2,109 genes that were DE in *arp6* (1,036 genes up-regulated and 1,073 genes down-regulated), 4,250 genes DE in *brm* (2,317 genes up and 1,933 genes down), and 3,203 genes DE in *arp6;brm* mutants (1,517 genes up and 1,686 genes down) (Fig. 1A, Summarized in Table S1). To determine the general processes influenced by ARP6 and BRM, we identified overrepresented gene ontology (GO) terms associated with all DE genes in each mutant compared to WT (Table S2).

**Figure 1.**
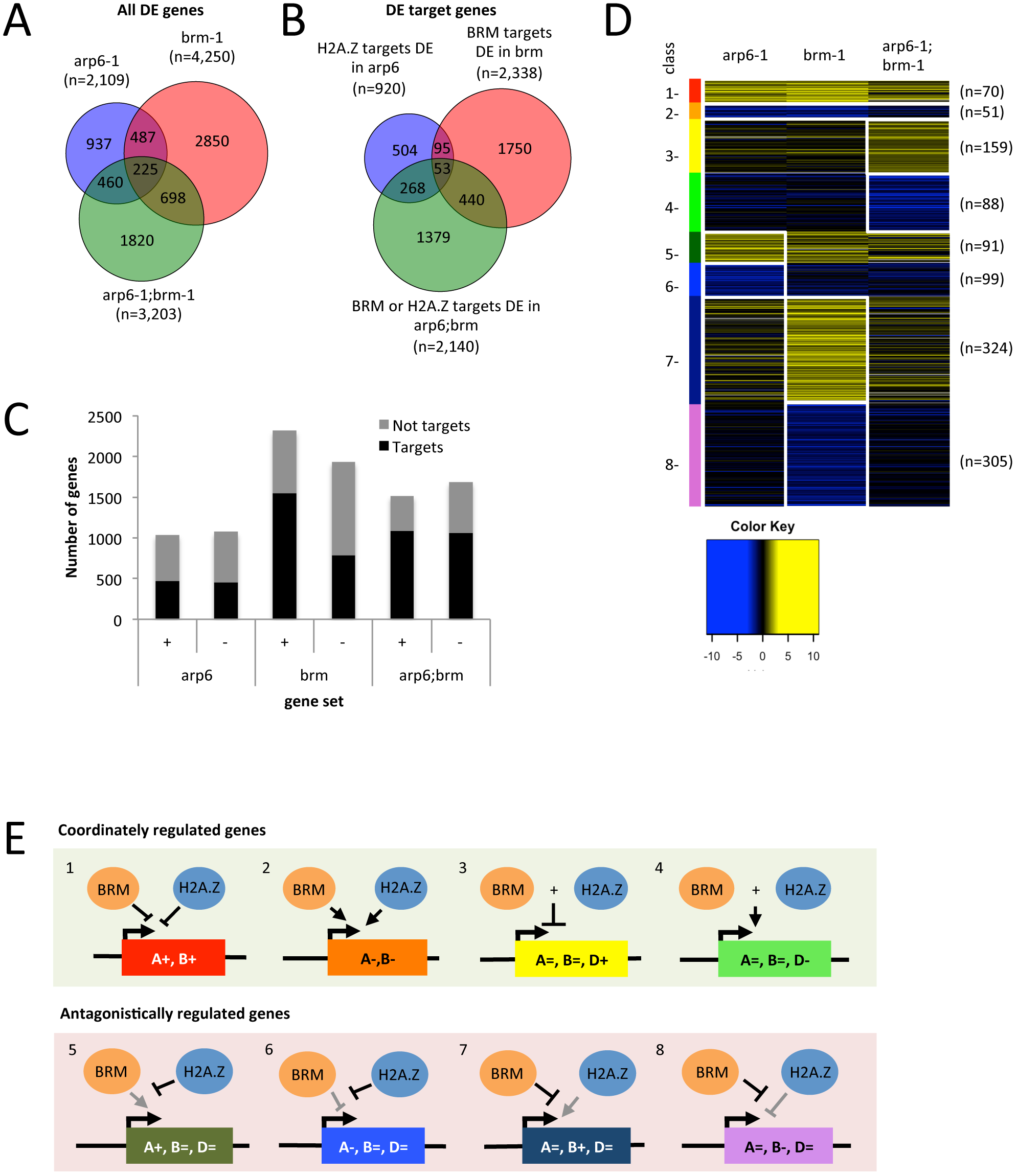
H2A.Z and BRM regulate transcription through various cooperative and antagonistic relationships. **(A)** Venn diagram shows the number of differentially expressed (DE) genes in each genotype and their significant overlap (hypergeometric test, p-value >0.001) with the genes DE in other mutants relative to WT. **(B)** Venn diagram shows the number of H2A.Z target genes DE in *arp6–1*, BRM target genes DE in *brm-1* mutants, and H2A.Z or BRM target genes DE in *arp6–1;brm-1* mutants relative to WT as well as their degree of overlap (hypergeometric test, p>0.001). **(C)** Histogram showing the number of direct DE target genes (black) and DE genes that are not targets (grey) that are up-regulated (+) or down-regulated (-) in *arp6, brm*, or *arp6;brm* mutants compared to WT. **(D)** Heatmap showing the average log_2_ fold change of genes in 8 classes of genes regulated antagonistically and coordinately by BRM and H2A.Z. Genes up-regulated are indicated in yellow and genes that are down-regulated are indicated in blue with a gradient black representing no change in transcription. The 8 different classes are indicated by various colors to the left and are divided into gene classes based on their pattern of transcription change in the different genotypes. Coordinately regulated genes include those that are up- (1-red) or down-regulated (2-orange) in both *arp6* and *brm* mutants relative to WT, genes that are up- (3-yellow) or down-regulated (4-light green) in the *arp6;brm* double mutant, but not the single mutants. Antagonistically regulated genes are divided into those genes that are up- (5-dark green) or down-regulated (6-light blue) in the *arp6* mutants but not *brm* or *arp6;brm* double mutants relative to WT, or genes that are up- (7-dark blue) or down-regulated (8-pink) in the *brm* mutants but not *arp6* or *arp6;brm* double mutants relative to WT. White boxes around gene sets highlight the significantly DE genes used to define the corresponding gene class, and the number of genes in each class **(n)** is shown to the right of the heatmap. **(E)** Diagram depicting the 8 genetic relationships between BRM and H2A.Z at coordinately regulated gene sets (top, green box) and antagonistically regulated gene sets (bottom, red box). The number to the top left of each diagram and the color of the targeted gene in the diagram correspond to the gene class defined in the heatmap **(D)**. The text within the target gene describes the transcription level relative to WT as increasing (+), decreasing (-), or not changing (=) in *arp6* **(A)**, *brm* **(B)**, or *arp6;brm* **(D)** mutants. Arrows represent a positive contribution to transcriptional regulation; lines with blunt ends represent a negative contribution to transcriptional regulation. Plus signs (+) between BRM and H2A.Z indicate a combined contribution to regulate transcription. Grey lines indicate the contribution of the opposing factor in the absence of the other factor.

In *Arabidopsis*, BRM regulates many developmental processes and responses to environmental stimuli, and integrates signals to allocate resources between stress response and growth (Farrona et al., 2011; Han et al., 2012; Wu et al., 2012;; Efroni et al., 2013; Archacki et al., 2013; Li et al., 2015a; Yang et al., 2015; Zhao et al., 2015; Archacki et al., 2016; Brzezinka et al., 2016; Zhang et al., 2016). H2A.Z also facilitates the transcriptional activation and repression of environmentally responsive genes (Coleman-Derr and Zilberman 2012; Berriri et al., 2016; Sura et al., 2017). In our data, we find a similar GO enrichment for DE genes in *arp6* and *brm* mutants related to responses to environmentally regulated genes, development, and transcriptional regulation (Fig. 2A-C). GO terms specifically overrepresented in the DE genes in *arp6* mutants include rRNA processing/modification, response to gibberellin stimulus, and defense response to virus (Fig. 2A and Table S2). GO terms specific to the DE genes in *brm* mutants include peptide transport, pollen-pistil interaction, cold acclimation, systemic acquired resistance, and translation, and some unique responses to hormones and abiotic stimuli (Fig. 2C and Table S2).

**Figure 2.**
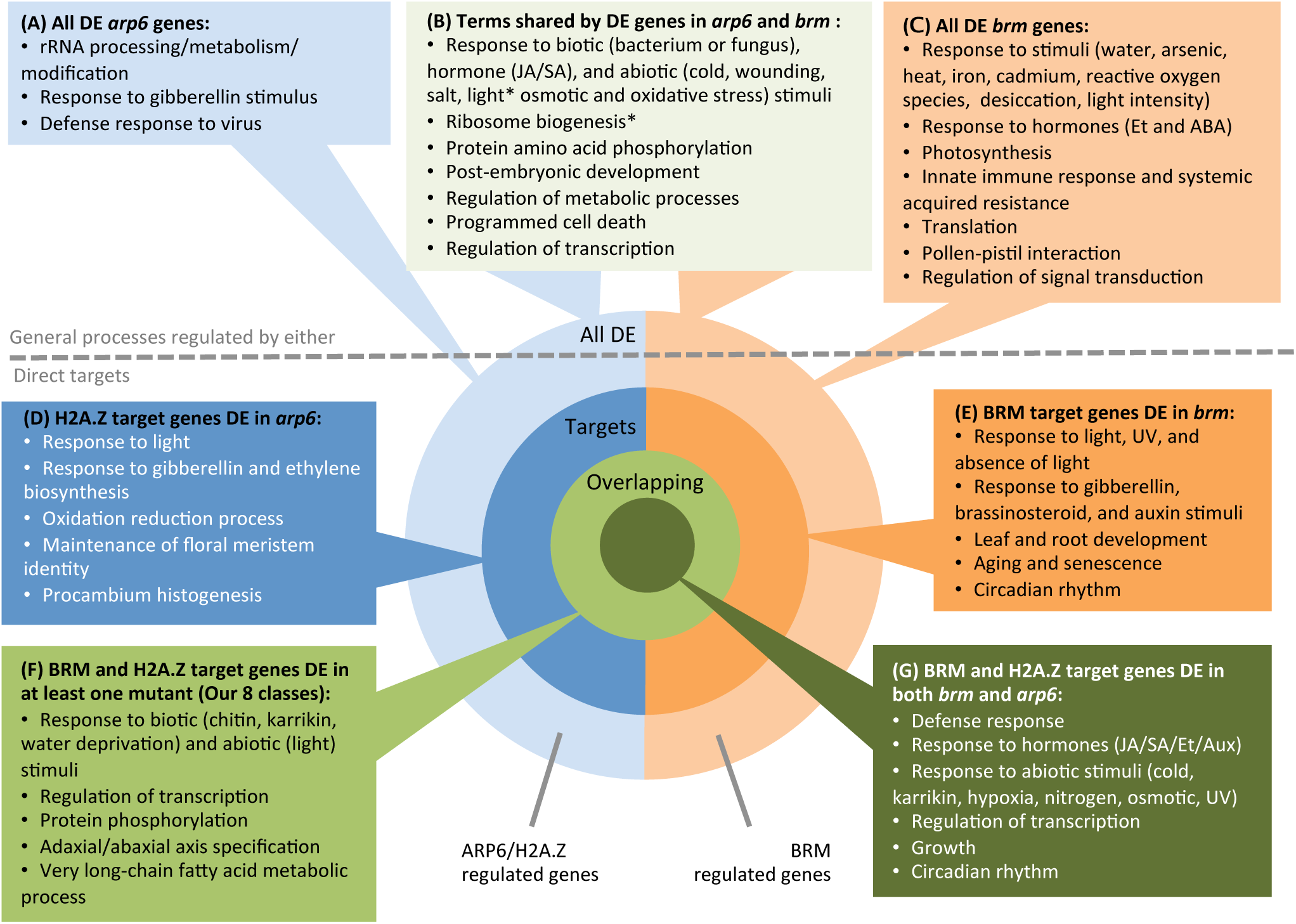
Gene Ontology (GO) analysis summary of BRM and ARP6/H2A.Z regulated genes. The circle displays the types of genes surveyed for GO analysis: all genes DE in *arp6* (A, light blue), all genes differentially expressed (DE) in *brm* (C, light orange), and the genes that are DE that are also targets of H2A.Z (D, dark blue) or BRM (D, dark orange), followed by genes that are DE targets of both H2A.Z and BRM and DE in at least one mutant (F, bright green), or both *arp6* and *brm* (G, dark green). Boxes surrounding the circle diagram list the terms associated with each category. Hormones are abbreviated as JA - jasmonic acid, ABA - abscisic acid, Aux-auxin, and Et - ethylene. The light green box **(B)** summarizes processes that are enriched in genes DE in both *arp6* and *brm* mutants and GO terms marked with an asterisk (*) represent terms that were enriched for the total set of DE genes in one mutant but that were enriched in the set of up- or down-regulated in the other mutant instead of the list of total DE genes. The grey dotted line separates the GO term summary boxes for general processes regulated by each factor from processes with genes that are direct targets of H2A.Z or BRM.

GO terms significantly overrepresented among DE gene sets from both *brm* and *arp6* mutants include responses to stimuli, regulation of transcription, and regulation of metabolic processes (Fig. 2B and Table S2). Stimuli highlighted by the analysis include external biotic stimuli such as response to bacterium or fungus, innate immune response, and defense response; endogenous stimuli such as hormone stimuli (jasmonic acid or salicylic acid); and abiotic stimuli such as cold, wounding, salt, osmotic stress, or oxidative stress, many of which have been reported previously (Coleman-Derr and Zilberman 2012; Berriri et al., 2016; Li et al., 2016; Archacki et al., 2016). Although many terms are shared between the lists of differentially expressed genes in *arp6* and *brm* mutants, it is worth noting that the actual genes that are associated with a shared term are not always the same (Table S2). This suggests that ARP6 and BRM contribute to similar general processes, albeit in some cases by affecting different genes.

### Assessing H2A.Z and BRM localization

Although many genes are DE in *arp6* and *brm* mutants, these are not necessarily direct targets of H2A.Z or BRM. Since ‘regulation of transcription’ is a GO term enriched in DE gene sets from each mutant, the gene products that are mis-expressed in the mutants may go on to cause secondary changes in transcription. To identify the genes directly targeted by H2A.Z and BRM, we analyzed our RNA-seq data in combination with data from BRM and H2A.Z chromatin immunoprecipitation experiments followed by sequencing (ChIP-seq). To assess BRM localization, we used previously published BRM-GFP ChIP-seq peaks generated from 14 day-old seedlings grown on plates (Li et al., 2016). Archacki et al. (2016) compared these ChIP-seq sites to their BRM ChIP-chip sites generated from 3-week-old plants grown on soil. The authors saw that even in different growth conditions and a different developmental stage, which consequently included different tissues, BRM was stably associated with similar sites with significant overlap between the two data sets (Archacki et al., 2016). Since our WT plants are at the same developmental stage as the published BRM ChIP-seq data we selected, we are confident that they sufficiently represent BRM localization for our experiments.

To assess H2A.Z localization relative to DE genes, we performed ChIP-seq for H2A.Z on green tissue from developmentally staged plants with 4–5 leaves from *arp6* mutants, *brm* mutants, and WT plants. Once sequence reads were mapped to the genome, Homer software (Heinz et al., 2010) was used to determine significant peaks in ChIP-seq read signal indicative of H2A.Z localization within each genotype. Since we are using *arp6* mutants as a proxy for H2A.Z mutants, we plotted H2A.Z ChIP-seq read signal from *arp6* mutants at H2A.Z peaks and confirmed that nucleosomes that normally contain H2A.Z are depleted of it in *arp6* mutants (Fig. S1A). To focus our analysis on sites where H2A.Z localization is dependent on ARP6, we removed the few H2A.Z peaks called in WT that overlapped with peaks called in *arp6* mutants (n=801). This left us with 11,877 ARP6-dependent H2A.Z peaks to assess how H2A.Z localization relates to transcriptional changes observed in the *arp6* mutants.

### Identifying differentially expressed H2A.Z and BRM target genes

To determine which of the transcriptional changes detected in our RNA-seq data are directly associated with H2A.Z and BRM localization, we identified DE target genes for either factor by integrating transcriptome data with H2A.Z and BRM ChIP-seq data. If an H2A.Z or BRM ChIP-seq peak fell within a gene body or if the closest TSS to a binding site was a DE gene, the gene was considered a target of that factor. Only 471 (45%) of up- and 449 (42%) of down-regulated *arp6* DE genes are directly associated with an ARP6-dependent H2A.Z peak (Fig. 1B and C and Table S1). In *brm* mutants, 1,552 (67%) of up- and 786 (41%) of down-regulated genes are direct targets of BRM (Fig. 1B and C and Table S1). In *arp6;brm* double mutants, the changes in gene expression were considered direct effects of the mutations if they were targets for H2A.Z, BRM, or both. Therefore, 1,082 (71.3%) were up- and 1,058 (62.8%) were down-regulated targets of either factor in the *arp6;brm* mutants (Fig. 1B and C and Table S1). Therefore, we defined H2A.Z and BRM DE target genes 1) by RNA-seq data showing that the genes are DE in the *arp6, brm*, or *arp6;brm* mutants respectively, as well as 2) using ChIP-seq data to confirm that H2A.Z or BRM are normally found at these genes in WT plants. From this point on in the study, we focused our analysis on the DE BRM and ARP6-dependent H2A.Z target genes as genes whose transcription is directly regulated by H2A.Z/ARP6 and BRM. Collectively, the defined DE target genes support the notion that both BRM and H2A.Z contribute to gene repression and activation in different contexts (Marques et al., 2010; Archacki et al., 2016).

### H2A.Z and BRM directly regulate transcription of developmental and environmental response genes

To better understand what types of genes are directly targeted by H2A.Z and BRM, we performed a GO analysis on DE genes in each mutant that are direct targets of both factors. We identified many interconnected terms for genes targeted by both H2A.Z and BRM that relate to development/growth and responses to environmental stimuli (esp. fungal response, osmotic stress, cold, and light) and hormones (auxin, ethylene, jasmonic acid, salicylic acid) (Fig 2F and G and Table S2). Demonstrating the interconnected nature of these terms, we know that the plant transcriptional network for responding to cold shares components with both defense and light stimuli responses (Catala et al., 2011; Barah et al., 2016). Also, the hormone response and metabolic process terms identified also relate to various aspects of defense response (Alazem and Lin 2015; Hiruma et al., 2010; Alazem and Lin 2015).

The DE genes that are direct targets of either H2A.Z in *arp6* mutants or BRM in *brm* mutants are individually enriched for similar GO terms as the list of GO terms that describe the total list of DE genes in either mutant. Of note, response to giberellin and red light were processes enriched in the DE target genes from either *arp6* or *brm* mutants individually, even though they were not processes that were overrepresented in the list of genes that are targets of both H2A.Z and BRM (Fig. 2D and e). Also, GO terms relating to aging and sensence were enriched in the list of BRM targets uniquely DE in *brm* mutants, but were not identified in the total list of DE genes and have not been reported before in BRM studies (Fig. 2E). Further work to understand whether any of these transcriptional changes produce phenotypes that are present in the *arp6;brm* double mutants, but not the single mutants will show whether BRM and H2A.Z have addititve, redundant, or antagonistic roles in these processes.

### H2A.Z and BRM coordinately and antagonistically regulate gene transcription

Since Farrona et al. (2011) suggested that H2A.Z and BRM antagonistically regulate transcription of the *FLC* gene in *Arabidopsis*, we wanted to test whether this antagonistic relationship extends to other genes across the genome. We also tested the hypothesis that H2A.Z and BRM could work together to regulate gene transcription as suggested by their roles in yeast (Santisteban et al., 2000). After verifying which differentially expressed genes are direct targets of H2A.Z and BRM using ChIP-seq data, we identified 8 gene classes that are either coordinately or antagonistically regulated by H2A.Z and BRM based on whether their transcript levels changed in one or more mutants (Fig.1 D and Table S1). To describe these gene classes, we will refer to genotypes and transcriptional changes with the following abbreviations: *A=arp6–1, B=brm-1, D=arp6–1;brm-1* double mutant, “+” = genes up-regulated in the specified mutant, “-” = genes down-regulated in the specified mutant, “=” = genes not DE in the specified mutant, n = number of genes in the class. We first identified genes that are coordinately regulated by H2A.Z/ARP6 and BRM. This category includes target genes that are up-regulated in both *arp6–1* and *brm-1* mutants compared to WT plants (Class 1: A+, B+, n=70), targets down-regulated in both *arp6* and *brm* mutants relative to WT (Class 2: A-, B-, n=51), and target genes with no change in transcript level in the individual mutants, but that are up- (Class 3: A=,B=,D+, n=159) or down-regulated (Class 4: A=, B=, D-, n=88) in the *arp6;brm* double mutants relative to WT (Fig. 1D and E). Classes 1 and 2 indicate that both H2A.Z and BRM are independently required for the proper regulation of these genes (Fig 1E). Classes 3 and 4 are DE target genes in the double mutants but not the single mutants, which are particularly interesting because these are genes where BRM and H2A.Z appear to work redundantly (Fig. 1E).

We also identified different classes of target genes where H2A.Z and BRM act antagonistically. This category of genes includes those either up- or down-regulated in a single mutant but that are neither DE in the other mutant nor the double mutant (Class 5: A+, B=, D=, n=91; Class 6: A-, B=, D=, n=99; Class 7: A=, B+, D=, n=324; Class 8: A=, B-, D=, n=305) (Fig. 1D and E). Since the loss of the second factor suppresses the change in transcript levels observed in the single mutant, H2A.Z and BRM seem to have opposing functions at these genes that become evident in the single mutants (Fig. 1E). Using class 5 as an example, these are H2A.Z and BRM target genes that have increased transcript levels in the *arp6* mutants when H2A.Z is depleted from nucleosomes (Fig. 1D). These same genes however are no longer significantly differentially expressed relative to WT when BRM is also depleted in the double mutants (Fig. 1D). Therefore, it seems that H2A.Z does not merely play a repressive role at these genes but does so by opposing the positive regulatory contribution of BRM at genes in class 5 (Fig. 1E). Alternatively, at genes in class 6, H2A.Z opposes the repressive role of BRM (Fig. 1E). Reciprocally, BRM also opposes the positive and negative contributions of H2A.Z to transcriptional regulation in classes 7 and 8, respectively. The mechanisms behind how H2A.Z and BRM positively and negatively regulate transcription at these genes, and how one opposes the function of the other still remain to be determined.

To determine the processes that may be influenced by each genetic interaction between H2A.Z and BRM, we evaluated GO terms for biological processes significantly enriched in our 8 coordinately or antagonistically regulated gene sets (Table S2). Genes in Class 4 (A=,B=,D-) do not have any significantly enriched GO terms, but the other classes were enriched for developmental and responsive processes (Table S2). Response to light stimulus is enriched in three gene classes (Class 6: A-, B=, D=; Class 7: A=,B+, D=; Class 8: A=,B-, D=), response to karrakin is enriched in 5 classes (Class 3: A=, B=, D+; Class 5: A+, B=, D=; Class 6: A-, B=, D=; Class 7: A=, B+, D=; Class 8: A=, B-, D=), and plant-pathogen interaction and defense response to fungus are enriched for three gene sets that include genes that are up-regulated in one or both genotypes (Class 1: A+, B+; Class 5: A+, B=, D=; Class 7: A=, B+, D=) (Table S2). Additionally, sequence-specific DNA binding transcription factor activity was enriched for 5 out of the 8 gene sets, emphasizing the roles of BRM and H2A.Z again in modulating other processes indirectly by controlling the expression of transcriptional regulators. The GO terms collectively enriched for genes in the 8 classes of H2A.Z and BRM co-targets DE in at least one mutant are similar to those target genes DE in both mutants (Fig. 2F and G and Table S2).

### H2A.Z localization is not dependent on BRM

One explanation for how BRM and H2A.Z/ARP6 could coordinately regulate gene expression (i.e. Figure 1E, Classes 1–4) is that BRM may regulate H2A.Z levels in chromatin. To assess whether BRM is important for H2A.Z occupancy, we analyzed H2A.Z levels at regions significantly enriched for H2A.Z in either WT plants or *brm* mutants. Some sites with significant H2A.Z localization in WT plants were not identified as significant in *brm* mutants based on peak calling alone, and reciprocally, some sites of enrichment were identified in *brm* mutants but not in WT plants (Fig. S1A). To see if sites unique to one genotype represent sites of H2A.Z depletion or gain in *brm* mutants, we plotted the H2A.Z ChIP-seq read signal from WT and *brm* plants across these regions of H2A.Z enrichment unique to either genotype. We observed comparable levels of H2A.Z enrichment in both genotypes even though the read signal did not meet the peak calling threshold for both genotypes (Fig. S1B). Since we only observed marginal differences between in H2A.Z levels in *brm* mutants and WT plants, this suggests that H2A.Z levels in chromatin are not generally dependent on BRM.

### BRM contributes to nucleosome stability and positioning at nucleosome depleted regions

The chromatin remodeling roles of BRM as a SWI2/SNF2 ATPase and H2A.Z incorporation into nucleosomes by the sWR1 CRC both can affect nucleosome stability and positioning at individual loci (Han 2012; Wu et al 2012; Brzezinka et al., 2016; Rudnizky et al., 2016). Identifying sites where both H2A.Z and BRM influence chromatin organization allows us to determine whether the presence of one could antagonize or enhance the chromatin modulating function of the other. To assess the genome wide contributions of H2A.Z and BRM to nucleosome organization, we performed Micrococcal Nuclease (MNase) digestion followed by sequencing (MNase-seq) on green tissue from 4–5 leaf developmentally-staged *arp6, brm*, and *arp6;brm* and WT plants. The nuclease activity of MNase specifically digests nucleosome-free DNA and leaves behind nucleosome-protected DNA, which provides a measure of where and how often a nucleosome is associated with a locus in our material (Allan et al., 2012). Thus, MNase-seq experiments allow us to evaluate how H2A.Z and BRM influence nucleosome occupancy and positioning (Allan et al., 2012; Zhang et al., 2015). Using H2A.Z and BRM ChIP-seq data, we evaluated nucleosomal changes that occur in our mutants at sites enriched for either H2A.Z, BRM, or both in order to focus our analysis and describe the ways that H2A.Z and BRM influence nucleosome stability and positioning.

To survey to what extent BRM contributes to nucleosome occupancy and positioning across the genome, we evaluated nucleosome dynamics in *brm* mutants compared to WT plants using MNase-seq data. By plotting nucleosome levels across sites where BRM localizes, we found that BRM is enriched at nucleosome-depleted regions (NDRs) and is often flanked on either side by well-positioned nucleosomes (Fig. 3A). Some of these well-positioned flanking nucleosomes become more stable in the absence of BRM in the *brm* mutant, suggesting that BRM contributes to nucleosome destabilization at the regions bordering where it binds. These results support previous findings that have shown another SWI2/SNF2 subunit also localizes to NDRs (Jegu et al., 2017). They also expand on the observation that BRM localizes between two well-positioned nucleosomes in a site-specific study since we show that finding two well-positioned nucleosomes flanking BRM sites is a genomic trend (Wu et al., 2015).

**Figure 3.**
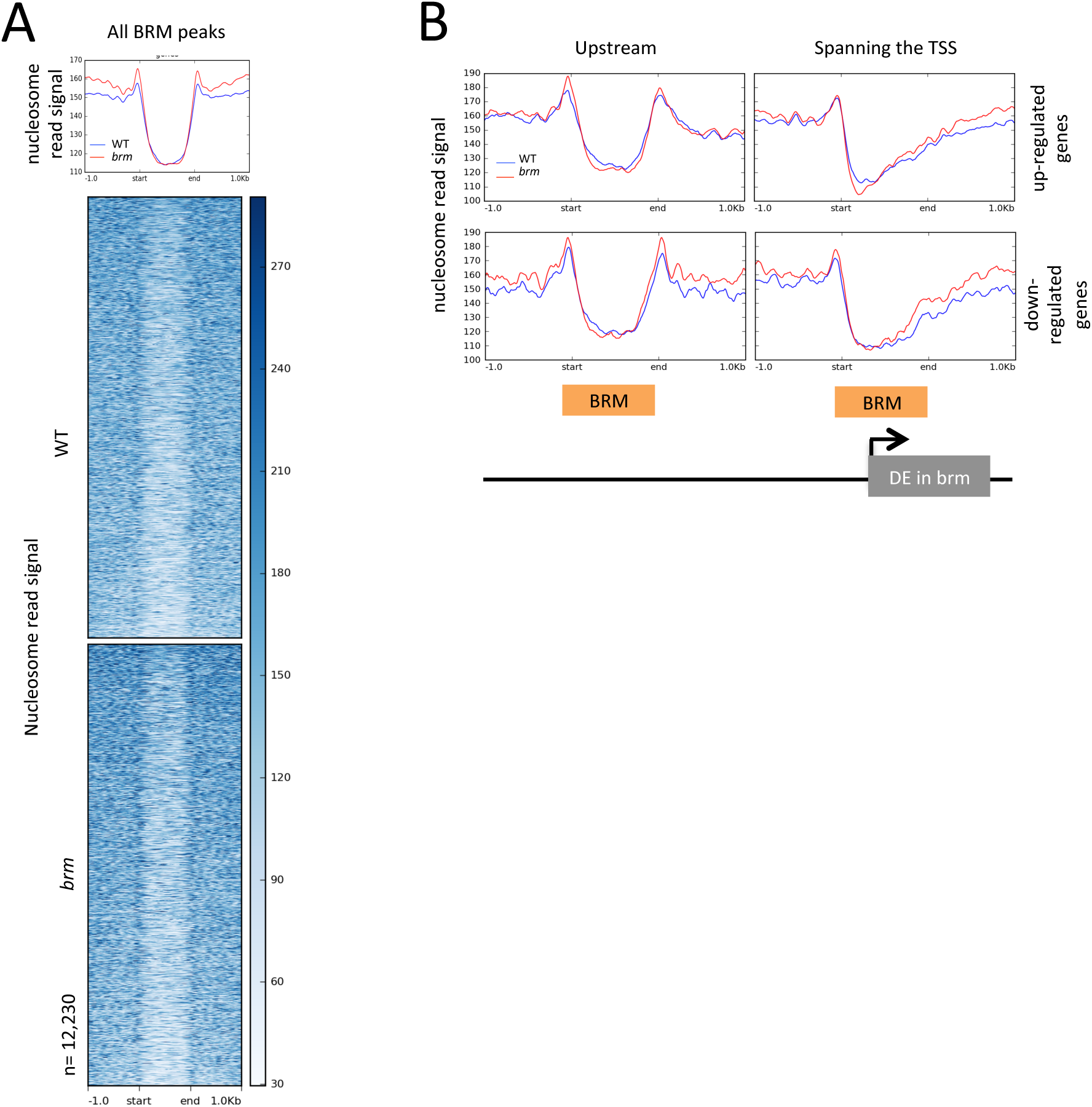
BRM is flanked by two well-positioned nucleosomes that are disrupted by transcription. **(A)** Profile plot and heatmap showing nucleosome read signals ± 1kb around all BRM peaks size-scaled to be 1 kb wide. Nucleosome reads are from an MNase-seq experiment in WT (blue line, top box of heatmap) and *brm* mutants (red line, bottom box of heatmap). **(B)** Average profile plots show nucleosome read signals from WT (blue) and *brm* (red) plants at select BRM ChIP-seq peaks associated with DE genes. Signals are plotted ± lkb around the start and ends of peaks scaled to be 1 kb in width. BRM peaks were divided into those **(i)** up-stream of or (ii) spanning the transcription start site (TSS) of genes up-regulated in *brm* mutants or BRM peaks that are (iii) up-stream of or (iv) span the TSS of a gene down-regulated *in brm* mutants. The diagram below the plots illustrates the general position of BRM peaks (orange box) used for the plots relative to the start of genes that are DE in *brm* mutants.

The well-positioned nucleosomes on either side of the BRM peaks could play a role in transcriptional regulation and be impacted specifically where we see transcriptional changes in *brm* mutants. To test whether these well-positioned nucleosomes are impacted by changes in transcription, we plotted the nucleosome signals from WT and *brm* mutants around BRM ChIP-seq peaks that were located either upstream of TSSs or spanning the TSSs of genes that are either up- or down-regulated in *brm* mutants relative to WT plants (Fig. 3B and Fig. S2A). When BRM localizes to the TSS of a gene that is either up- or down-regulated, one well-positioned nucleosome was found upstream of the TSS, but to the downstream side of the BRM ChIP-seq peak DNA was more accessible to MNase digestion (Fig. 3B). One explanation for this is that BRM localization actually extends past the defined BRM peaks. However, by plotting BRM ChIP-seq signal at defined BRM ChIP peaks, we find that the more open chromatin conformation extends past BRM enriched sites in the direction of transcription (Fig. S2B). Another explanation is that other factors may override the contribution BRM makes to nucleosome positioning downstream of BRM peaks, so that there is a more dispersed nucleosome signal relative to BRM binding when genes are transcribed. Based on the nucleosome average plot profiles, there were no significant changes in nucleosome occupancy at BRM peaks at either up- or down-regulated genes in *brm* mutants compared to WT, although there was some increase in nucleosome occupancy directly downstream of where BRM localized (Fig. 3B). When BRM is found upstream of the TSS, it is still flanked by well-positioned nucleosomes on both sides (Fig 3B). K-means clustering did not indicate that there were any subsets of upstream BRM peaks that might show the same degree of directionality that was observed for peaks that associate with TSSs (Fig. S2C). Together, the nucleosome patterns at BRM peaks support the notion that BRM binds both to the NDR adjacent to the TSS and also to upstream sites with open chromatin structure, such as regulatory regions within promoters or enhancers. We do not observe notable changes in nucleosome occupancy across all sites where BRM localizes when we compare between WT and *brm* mutant nucleosomes (Fig. 3A). Therefore, in a general sense it seems that BRM is not required to produce these NDRs but may perform other functions once targeted there.

Since locus specific studies have described roles for BRM and other SWI2/SNF2 subunits in nucleosome positioning and destabilization, we decided to quantify how often BRM is associated with different types of nucleosome dynamics (Han et al 2012; Wu et al., 2015; Brzezinka et al., 2016; Sacharowski et al., 2015). Using DANPOS2 software, nucleosomes were defined as dynamic if they had significant changes in nucleosome positioning (different position of nucleosome read summits), occupancy (different height of nucleosome read summits), fuzziness (difference in the standard deviation of nucleosome read positions), or any combination of the three in mutants relative to nucleosomes in WT tissue (Chen et al., 2013). We found that 25% of BRM peaks have significant nucleosome changes in *brm* mutants (based on an FDR cutoff of <0.05) (Fig. S3A). The chromatin landscape flanking BRM binding sites appears to have different nucleosome occupancy levels in *brm* mutants than those within BRM binding sites, based on the previous nucleosome read plots (Fig 3A). Therefore, we evaluated whether the types of nucleosome changes at the bordering regions of BRM ChIP-seq peaks are enriched for different types of changes than what is observed for nucleosomes found where BRM directly associates within peak centers (illustrated in Fig. 4A). For this purpose, we defined BRM peak borders as a ± 200 bp range around the start or end of a BRM ChIP-seq peak to describe how BRM contributes to nucleosome dynamics at the well-positioned nucleosomes that flank BRM sites (Fig. 4A). We further separated peaks into size quartiles so that the largest and smallest peaks would not skew observations at intermediate sized peaks. This also allows for a more relative comparison between changes observed within the standard 400 bp sized border regions we defined and the BRM ChIP-seq peak centers, which have variable sizes (from 300 bp to 4kb).

**Figure 4.**
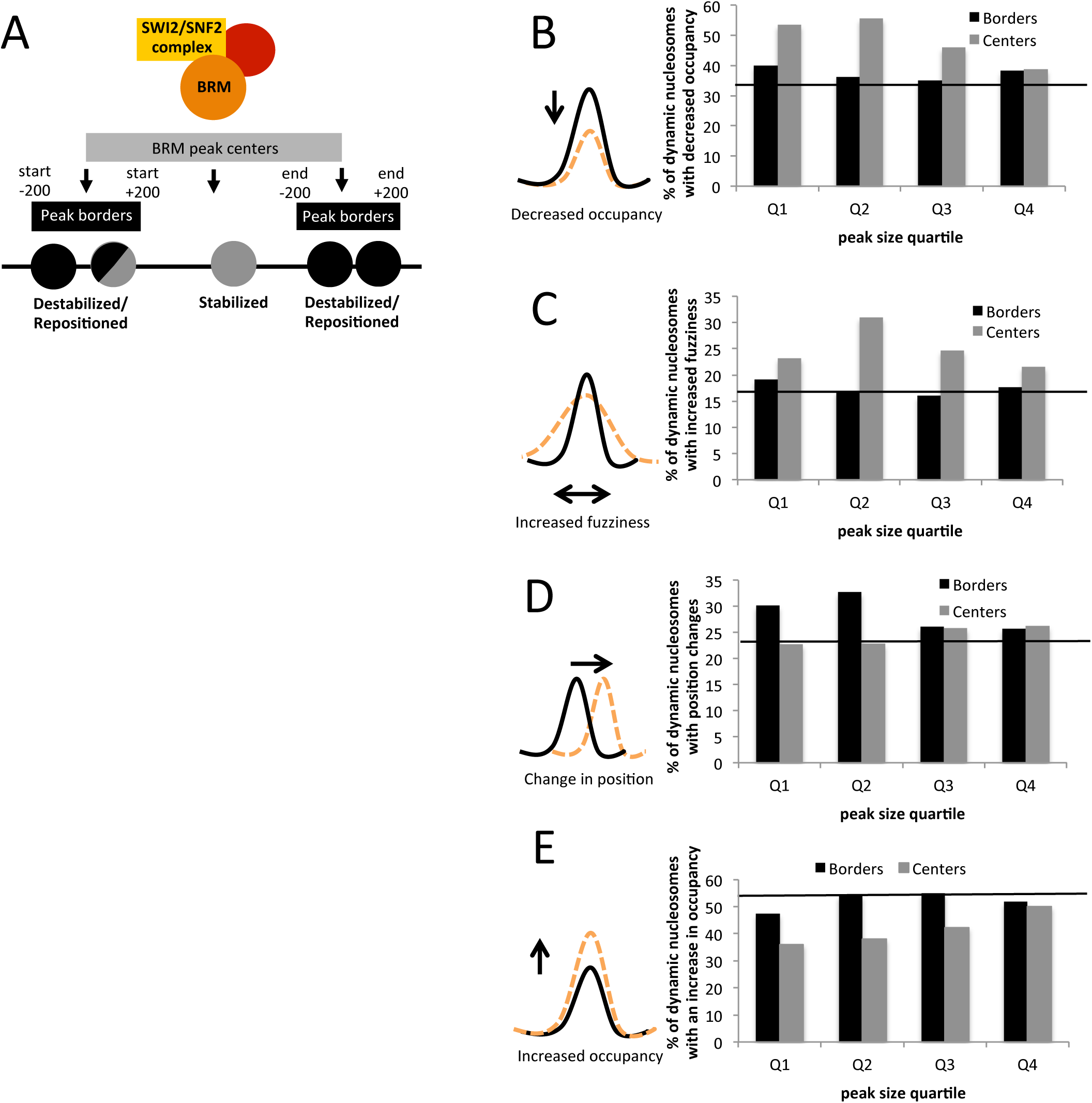
BRM contributes to nucleosome stability and positioning differentially at nucleosome depleted regions and flanking areas. **(A)** Depiction of BRM in *the Arabidopsis* SWI2/SNF2 complex associating with chromatin and stabilizing nucleosomes within peaks and repositioning nucleosomes at peak borders. Nucleosomes within the center are colored grey and nucleosomes within border regions are colored black. Two-toned nucleosomes represent how the data are not mutually exclusive but can contain nucleosomes that fall in both border and center categories. **B-E**) Histograms summarizing the proportion of dynamic nucleosomes found at the borders (black) or centers (gray) of BRM peaks that show a decrease in occupancy **(B)**, change in position **(C)**, an increase in fuzziness **(D)**, or an increase in occupancy **(E)** in *brm* mutants. Peak centers included nucleosomes fully contained within the defined peaks and borders include those that overlap with a peak start or end. The proportion of dynamic nucleosomes across the genome with the indicated type of change in *brm* mutants is shown as a black line for comparison. BRM peaks were separated into size quartiles for analysis: Q1: 300–415 bp, Q2: 415–546 bp, Q3: 546–776 bp, Q4: 776 bp-4kb. Diagrams to the left of the histograms represent the nucleosome changes described in *brm* mutants (orange dashed line) compared to WT nucleosomes (black line).

To identify how often nucleosome occupancy, positioning, or fuzziness depends on BRM, we quantified the proportion of different types of nucleosome changes that are observed among all nucleosomes considered dynamic between WT plants and *brm* mutants. Evaluating the *brm* mutant epigenome as a whole, 35% of the nucleosomes that change in *brm* mutants experience a decrease in occupancy. However, specifically evaluating changes that take place where BRM localizes directly (peak centers, and particularly the smaller ones in Q1 & Q2), 55% of the dynamic nucleosomes show decreases in occupancy (Fig. 4B). Similarly, 15% of the nucleosomes that change across the genome experience an increase in fuzziness in *brm* mutants, while 25–30% of the nucleosomes in smaller BRM peak centers experience an increase in fuzziness (Fig. 4C). This enrichment for decreased nucleosome occupancy in combination with enrichment for increased nucleosome fuzziness at BRM peaks in *brm* mutants suggests that BRM contributes to the stability of any nucleosomes that are found where BRM binds (Fig. 4A).

Next, we measured the types of nucleosomal changes that occur in *brm* mutants in the regions flanking BRM peaks in comparison to the types of changes directly where BRM binds. The proportion of dynamic nucleosomes that have increases in fuzziness and decreases in occupancy at BRM peak borders is comparable to the levels seen across the genome (Fig. 4B,C). However, the nucleosomes at BRM peak borders at lower quartiles show a greater proportion of nucleosome position changes (30–33%) relative to the proportion observed across the genome (25%) or at BRM peak centers (approx. 25%) (Fig. 4D). Of the nucleosomes that change at BRM peak borders in *brm* mutants, a greater proportion experience increases in occupancy than decreases in occupancy or changes in positioning or fuzziness (Fig. 4B-E). Therefore, our MNase-seq data in combination with BRM ChIP-seq data demonstrate that BRM localizes to NDRs and contributes more to nucleosome stability where it directly associates with chromatin, while contributing more to destabilization or the positioning of flanking nucleosomes (Fig. 4A).

### H2A.Z has a variable influence on the surrounding nucleosome landscape

When H2A.Z is incorporated into nucleosomes, it can change both intra-nucleosomal interactions as well as the interactions between nucleosomes and other nuclear proteins (Bonisch and Hake 2012). Consequently, H2A.Z-containing nucleosomes have been associated with a range of nucleosome dynamics including changes in nucleosome stability and positioning (Bonisch and Hake 2012; Rudnizky et al., 2016). Before assessing whether specific types of nucleosomal changes are enriched at sites where H2A.Z is found in relation to BRM function, we used MNase-seq experiments to evaluate whether H2A.Z-containing nucleosomes are enriched for specific types of nucleosomal changes in *arp6* mutants compared to WT plants. By using MNase-seq data in combination with our H2A.Z ChIP-seq data, we evaluated whether specific types of nucleosomal changes (changes in positioning, occupancy, or fuzziness) were enriched at nucleosomes that normally contain H2A.Z, but that are depleted of H2A.Z in *arp6* mutants. Only a fraction of nucleosomes that normally contain H2A.Z in WT plants (14.6%, Fig. S3B) had significant changes when comparing nucleosomes in *arp6* plants using the DANpOS2 software. A similar, yet slightly higher proportion of the H2A.Z sites associated with up- or down-regulated genes in the *arp6* mutants contained significant nucleosome changes of at least one type, considering that 17.5% of H2A.Z peaks at genes down-regulated and 18.3% of H2A.Z peaks at gene up-regulated have significant nucleosome changes (Fig. S3B).

When we evaluated nucleosome occupancy in the *arp6* mutants, we noticed there were large gaps in nucleosome read signal (Fig. S4). We then compared our *arp6* MNase-seq nucleosome signals to *arp6* genomic DNA and found that the gaps in nucleosome read signal correspond to large genomic deletions in the *arp6* mutants (Fig. S4). These deletions would skew our MNase-seq results, making them appear as a loss of a nucleosome in the mutant compared to WT when instead there was a loss of genomic DNA in this mutant line. We therefore mapped the mutations using CNVnator software which reported 1,545 deletions (>200 bp) compared to the TAIR10 reference genome (Abyzov et al., 2011). Although some of these deletions are strain differences since they were also missing in our WT plants compared to the reference genome, the total deleted portion collectively covers 5.88 megabases of DNA, which is a large portion of the ~145 megabase Arabidopsis genome (Bevan and Walsh 2005). To ensure that we are analyzing nucleosome dynamics at regions of the genome that are present in *arp6* and *arp6;brm* mutants, we removed nucleosomes from our analysis if they were called as dynamic by DANPOS2 but also overlapped with deleted regions. We also required a minimum of 1 read per 10 bp area visualized as a cutoff when analyzing nucleosome plot profiles to exclude missing regions from our analyses.

After accounting for the deleted regions, we quantified the proportion of nucleosomes that changed in *arp6* mutants that had changes in occupancy, fuzziness, or positioning, using the same definitions we used to analyze nucleosomal changes in *brm* mutants. Using H2A.Z ChIP-seq peaks to focus our analysis, we found that the collection of nucleosomes that normally contain H2A.Z in WT but lose H2A.Z in *arp6* mutants experience both increases and decreases in fuzziness, increases and decreases in occupancy, and changes in positioning (these categories are not mutually exclusive) (Fig. S5A-C and Fig. 5). We further evaluated whether H2A.Z-containing nucleosomes are enriched for any particular type of nucleosomal change in comparison to the levels that are observed across the genome. Even though H2A.Z has comparable levels of nucleosomes that experience increases and decreases in fuzziness upon H2A.Z depletion in *arp6* mutants (Fig. 5), the proportion of nucleosomes that become less fuzzy in *arp6* mutants at H2A.Z peaks (27.4%) is greater than what is observed across the genome (19.9%) (Fig S5C). This emphasizes the role that H2A.Z plays in nucleosome destabilization in the genome. There were no other enrichments of one type of nucleosomal changes where H2A.Z localizes compared to changes observed in the *arp6* genome as a whole. We also specifically evaluated the types of nucleosomal changes that are enriched at subsets of H2A.Z-containing nucleosomes where H2A.Z either contributes to transcriptional repression or activation. We observed enrichment for nucleosome position changes and a shift toward nucleosome destabilization (a depletion of less fuzziness and more with increased fuzziness) at genes that are up-regulated in the *arp6* mutants (Fig. S5A and C). This trend suggests that the role of H2A.Z in transcriptional repression has a greater correlation with nucleosome stabilization (decreasing fuzziness and inhibiting position shifts) than with nucleosome destabilization. However, this correlation is consistent with the types of changes we would expect to correspond with an increase in transcription and may not be directly due to H2A.Z function.

**Figure 5.**
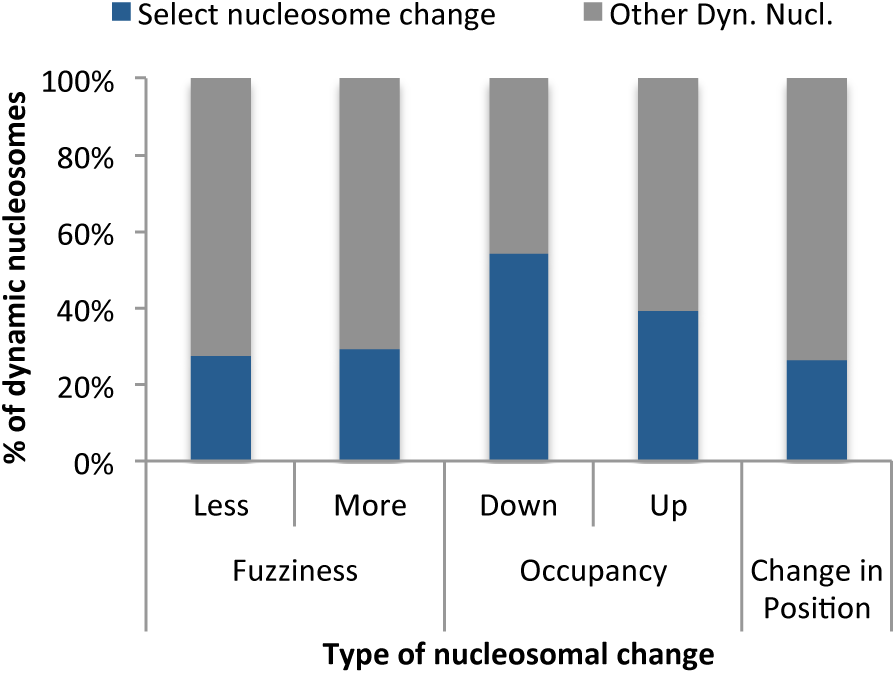
H2A.Z contributes to a range of nucleosome changes. Histogram summarizing the percentage of each type of DANPOS2-called dynamic nucleosomes (blue) compared to all other dynamic nucleosomes (grey) called within H2A.Z peaks when comparing *arp6* nucleosomes to WT nucleosomes.

When assessing the types of changes that coincide with H2A.Z depletion from nucleosomes that normally would contain it, it is important to compare them to changes observed at other places in the genome. Only 9.38% of total dynamic nucleosomes called between *arp6* mutants and WT are found where H2A.Z localizes (total dynamic nucleosomes=21,967, those at H2A.Z peaks=2,061). This means that other nucleosome changes are taking place in the genome due to secondary effects of H2A.Z depletion and SWR1 defects rather than the direct loss of H2A.Z alone. The preference for nucleosome occupancy changes in *arp6* dynamic nucleosomes may be attributable to changes in transcription, changes in chromatin organization/localization within the nucleus, or the direct effects of depleting chromatin of H2AZ alone. However, some secondary changes could also be due to some unaccounted deletions in the *arp6* genome.

### BRM contributes to nucleosome destabilization when it is in proximity to H2A.Z

Since both BRM and H2A.Z contribute to nucleosome stability and positioning individually, we wanted to evaluate whether they work coordinately or antagonistically on nucleosomes where they overlap. We defined 2,963 regions of overlap between BRM ChIP-seq peaks and H2A.Z ChIP-seq peaks (significant by Fisher’s exact test, p-value < 2.2e-16) (Fig. 6A). By plotting the ChIP-seq read signal for BRM and H2A.Z at these regions of overlap and dividing them into 4 K-means clusters, we determined that these are primarily regions of peripheral overlap instead of sites with strong co-localization (Fig. 6B). We then identified 88 regions of H2A.Z-BRM overlap that also contained significant nucleosome changes in both respective mutants compared to WT nucleosomes. These regions allow a more direct comparison between the roles that BRM and H2A.Z play in nucleosome dynamics. At these regions of shared overlap, H2A.Z and BRM both contribute to significant changes in nucleosome positioning at 42% (37/88) of nucleosomes, however only 21% (19/88) of these nucleosomes are changed in both genotypes. H2A.Z contributes evenly to increases and decreases in the degree of nucleosome occupancy changes (Fig 6C) and fuzziness changes (Fig 6D) in regions of BRM/H2A.Z overlap. These observations demonstrate that H2A.Z has a range of contributions to nucleosome stability at these sites, consistent with what is observed at H2A.Z sites alone (Fig. 5).

**Figure 6.**
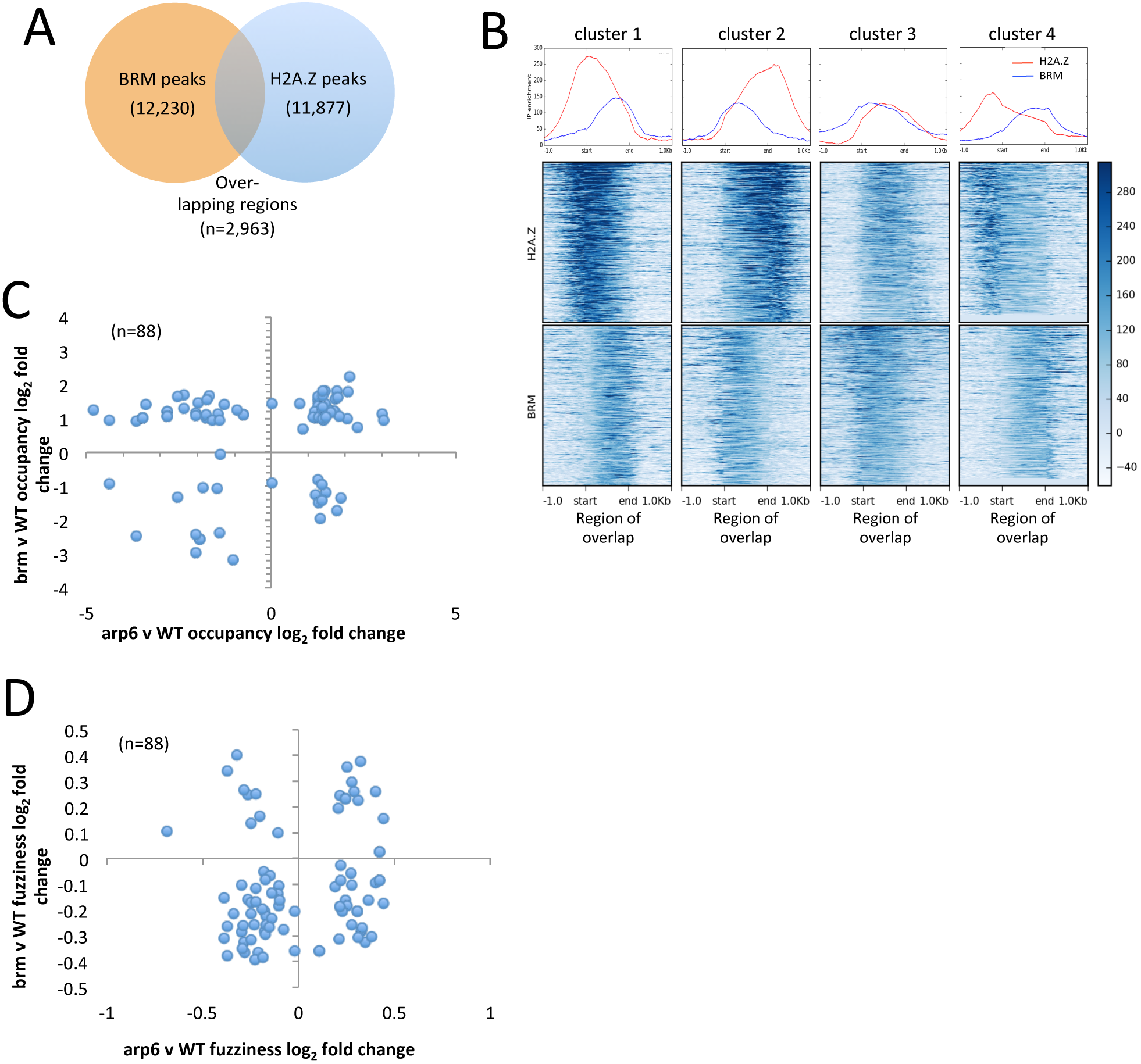
BRM destabilizes nucleosomes where BRM and H2A.Z overlap. **(A)** Venn diagram shows the number of BRM peaks, H2A.Z peaks, and regions of overlap between the two. **(B)** Average profile plots and heatmaps show four K-means clustered H2A.Z and BRM ChIP-seq read signal patterns (normalized to input) at regions of overlap between BRM and H2A.Z peaks. Regions of overlap with dynamic nucleosomes identified in each mutant relative to wild type nucleosomes (n=88) were further evaluated. Scatter plots display the log_2_ fold change in nucleosome occupancy **(C)** and fuzziness **(D)** in regions of H2A.Z/BRM overlap that contain dynamic nucleosomes in both *brm* mutants (y-axis) and *orp6* mutants (x-axis) compared to WT.

Alternatively, *brm* mutants have a greater proportion of nucleosomes with an increase in occupancy and decrease in fuzziness compared to WT at regions of H2A.Z/BRM overlap. These data indicate that BRM plays a greater role in nucleosome destabilization at sites where it overlaps with H2A.Z (Fig. 6C and D). This is consistent with the fact that there are more increases in nucleosome occupancy at BRM peak borders than at the centers (Fig. 3A and Fig. 4E) and that H2A.Z and BRM have more peripheral overlaps (Fig 6B). Collectively, these observations indicate that H2A.Z and BRM do not solely antagonize the function of the other in chromatin, but can also cause similar changes in nucleosome organization.

### BRM contributes to nucleosome destabilization of +1 nucleosomes at genes coordinately regulated with H2A.Z

To assess the roles of H2A.Z and BRM in nucleosome stability as they relate to transcriptional regulation, we plotted the average profiles of nucleosome read signals from WT, *arp6, brm*, and *arp6;brm* plants surrounding the transcription start sites (TSS) of DE genes (Fig. 7 and Fig. S6). We focused our analysis on TSSs from the 8 antagonistically or coordinately regulated DE H2A.Z/BRM target gene classes we identified earlier in the study based on their transcriptional changes in the mutants (Fig. 1C). While there are some nucleosome occupancy changes detected within gene bodies in mutants, we primarily focused our analysis on changes observed for the +1 nucleosome because it acts as a first physical barrier for transcriptional regulation (Weber et al., 2014).

**Figure 7:**
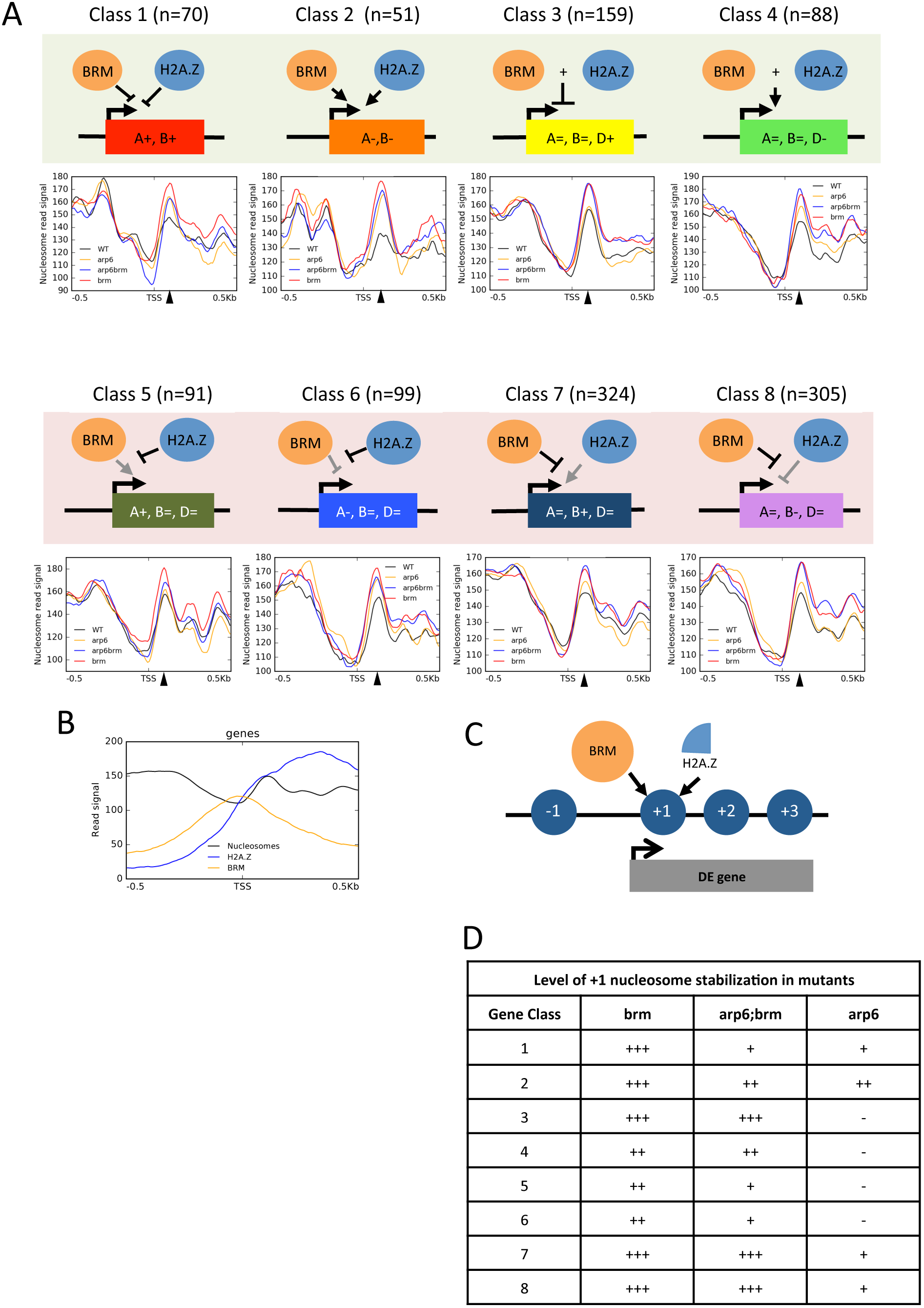
BRM and H2A.Z destabilize the +1 nucleosome at DE targets. **(A)** Profile plots showing the average nucleosome read signal from WT plants (black), *arp6* (orange), *arp6;brm* (blue), and *brm* mutants (red) ±500 bp around the TSSs of the 8 classes of DE H2A.Z and BRM target genes. Black triangles on the x-axis indicate the position of the +1 nucleosome. The diagrams above the plots describe the genetic relationships between BRM and H2A.Z/ARP6 for each gene class and are the same as those described in Fig. 1. **(B)** Profile plot showing the read signal for WT nucleosomes (black), H2A.Z ChIP-seq (blue), and BRM ChIP-seq (orange), averaged across ±500 bp up- and downstream of the TSSs of the DE BRM and H2A.Z target genes. **(C)** Diagram representing how we used ChIP-seq, MNase-seq, and RNA-seq data sets in the previous figures, to evaluated the relationship between BRM localization in WT (orange), H2A.Z localization in WT (light blue) and nucleosomes (blue circles) around the TSSs of DE BRM and H2A.Z target genes. **(D)** Table summarizes the extent to which the +1 nucleosomes become stabilized in *brm, arp6;brm* and *arp6* mutants at the 8 DE BRM-H2A.Z target gene classes defined in Fig 1. The level of nucleosome stabilization in the mutant was defined based on the overlaps between different measures of variance at the +1 nucleosome read signals plotted in Fig. S6. The degree to which the +1 nucleosome was stabilized in the mutants compared to WT is defined as no change (- = mean of one falls within the 95% confidence interval of the other); a small change (+ = the mean of one sample does not overlap with the 95% confidence interval of the other); a medium change (= the standard error of one sample does not overlap with the 95% confidence interval of the other); or a large change in nucleosome occupancy (+ = there is no overlap between 95% confidence intervals for the two samples).

At these DE gene classes, *brm, arp6* and *arp6;brm* mutants showed an increase in +1 nucleosome occupancy, with the most dramatic changes seen in the coordinately regulated gene Classes 1 and 2 (Fig. 7A and Fig. S6). The *brm* and *arp6;brm* mutants also show +1 nucleosome occupancy increases at gene Classes 3 and 4 (Fig. 7A and Fig. S6). It is interesting to note that the role of BRM in +1 nucleosome stabilization is unaffected by the direction of transcriptional change (Fig. 7A and D and Fig. S6). In *arp6* mutants, +1 nucleosome occupancy is mostly unchanged at genes DE in *arp6* or *arp6;brm* mutants (Classes 3–6) but show slight increases in nucleosome stability where H2A.Z opposes the regulatory functions of BRM at DE genes in *brm* mutants (Classes 7 and 8; Fig. 7A and D and Fig. S6). Although the loss of BRM results in +1 nucleosome occupancy increases at genes, especially in Class 3 and 4, significant changes in transcription do not happen in these gene classes until there is a loss of both H2A.Z and BRM in the *arp6;brm* mutants (Fig. 7A and Fig. S6). This means that at these gene classes (3 and 4), the increase in nucleosome stability in the BRM mutants is not sufficient to cause significant changes in transcription until H2A.Z is also lost.

BRM is enriched at the NDR just upstream of the +1 nucleosome at our 8 gene classes (Fig. 7B), so it may be influencing +1 nucleosome stability by interacting with the +1 nucleosome either peripherally or through recruiting other chromatin modifying factors to interact with the +1 nucleosome. Having more stable +1 nucleosomes at BRM targets in *brm* mutants is consistent with our observations that BRM contributes to nucleosome destabilization at the borders/flanking regions where it localizes and particularly when it co-localizes with H2A.Z (Fig 4C and E and Fig 6C and D). Further work to determine whether BRM destabilizes the +1 nucleosomes or whether it recruits or blocks other factors which indirectly contribute to +1 nucleosome stabilization will help us better understand the role of BRM in transcriptional regulation.

### BRM and H2A.Z may interact with TFs to facilitate transcriptional regulation

We originally wanted to expand on the work of Farrona et al (2011) to test the extent of the antagonistic relationship between BRM and ARP6/H2A.Z beyond what was observed at the *FLC* gene. Our work presenting variable nucleosome changes where the two factors overlap as well as at DE co-targeted genes (Fig. 6 and 7) indicates that the BRM-H2A.Z relationship is much more complex than a simple antagonism. H2A.Z and/or BRM may have more consistent roles in chromatin regulation as they relate to the functions of specific transcription factors (TFs). For example, both H2A.Z and the SWI2/SNF2 complex have been implicated in regulating chromatin accessibility for TFs (John et al., 2008; Sacharowski et al., 2015; Jegu et al., 2017). H2A.Z eviction from +1 nucleosomes is regulated by the HSFA1a TFs to regulate heat response genes (Cortijo et al., 2017) and conversely BRM can be recruited to chromatin by TFs (Wu 2012; Efroni et al., 2013; Vercruyssen et al., 2014; Zhao et al., 2015; Buszewicz et al., 2016; Zhang et al., 2016). These previously defined relationships between H2A.Z, BRM, and TFs prompted us to evaluate how H2A.Z and BRM contribute to nucleosome organization surrounding TF binding sites where they co-localize.

To identify TFs that may be associated with specific regulatory relationships between H2A.Z and BRM, we identified significantly enriched sequence motifs found in accessible chromatin regions associated with the 8 DE gene classes we identified (Fig 1D and E). Accessible chromatin sites were defined in a previous study using an ATAC-seq data set from leaf mesophyll cells, which is the predominant cell-type in our tissue (Sijacic et al., 2017). The motifs enriched at accessible regions across 7 of our DE gene classes are statistically similar to the target motifs for 78 different TFs (none were enriched for gene class 1) (Table S3). Of the factors identified, 15 have previously been reported to associate with the SWI2/SNF2 complex in Arabidopsis (Table S3; Efroni et al., 2013, Jegu et al., 2017; Zhang et al., 2017a).

Several of the TFs identified are involved in responses to light (SOC1, FRS9, HY5, MYC2, CIB2, BZR1, BIM1/2/3, and PIF1/3/4/5/7). These factors are intriguing because they are consistent with the multiple GO terms relating to responses to light stimuli that are enriched in our 8 classes of DE H2A.Z-BRM target genes (Figure 2 and Table S2). Both H2A.Z and BRM also independently regulate genes in response to various light stimuli (Table S2).

One family of light responsive TFs that was predicted to associate with both coordinately-regulated and antagonistically-regulated gene classes is the basic helix-loop-helix, PHYTOCHROME-lNTERACTlNG FACTOR (PIF) family of transcription factors (PIF1, 3, 4, 5, and 7) (Table S3). These TFs act as both positive and negative regulators of transcription, similar to H2A.Z and BRM (De Lucas and Prat 2014; Lee and Choi 2017). PIFs also integrate light response and hormone signals in plants to regulate growth and development in response to changes in light stimuli similar to the types of genes that are DE targets of H2A.Z and BRM (Fig 2) (De Lucas and Prat 2014; Lee and Choi 2017). Relationships between PIF TFs and either H2A.Z or the SWI2/SNF2 complex have been described before, making the PIF TFs interesting candidates for follow-up analyses (Efroni et al., 2013; Wigge 2013; Galvao et al., 2015; Jegu et al., 2017; Zhang et al., 2017a).

Before testing whether BRM or H2A.Z affect chromatin organization at PIF TF binding sites, we assessed the degree of overlap between BRM, H2A.Z and previously reported binding sites for PIF4 and PIF5 (Pedmale et al., 2016). We found that PIF4 and PIF5 peaks overlap with a combination of H2A.Z and BRM together, H2A.Z only, BRM only, or neither, with a slight preference toward overlapping with BRM rather than H2A.Z (Fig. 8A, Fig. S7A). To evaluate H2A.Z and BRM localization relative to PIF4 sites that overlap with H2A.Z and BRM, we plotted H2A.Z and BRM ChIP-seq signals (normalized to input) from WT plants relative to WT nucleosome patterns across the PIF4 ChIP-seq peaks (Fig 8B). Using K-means clustering to separate the nucleosome profiles around PIF4 sites into four different subsets, we found that BRM is enriched at the center of PIF4 peaks and H2A.Z is enriched at one or both sides of the PIF4 peaks (Fig. 8B). This suggests that BRM preferentially interacts with PIF4 binding sites and H2A.Z has a more peripheral interaction, consistent with BRM binding to NDR and H2A.Z localizing within flanking nucleosomes.

**Figure 8.**
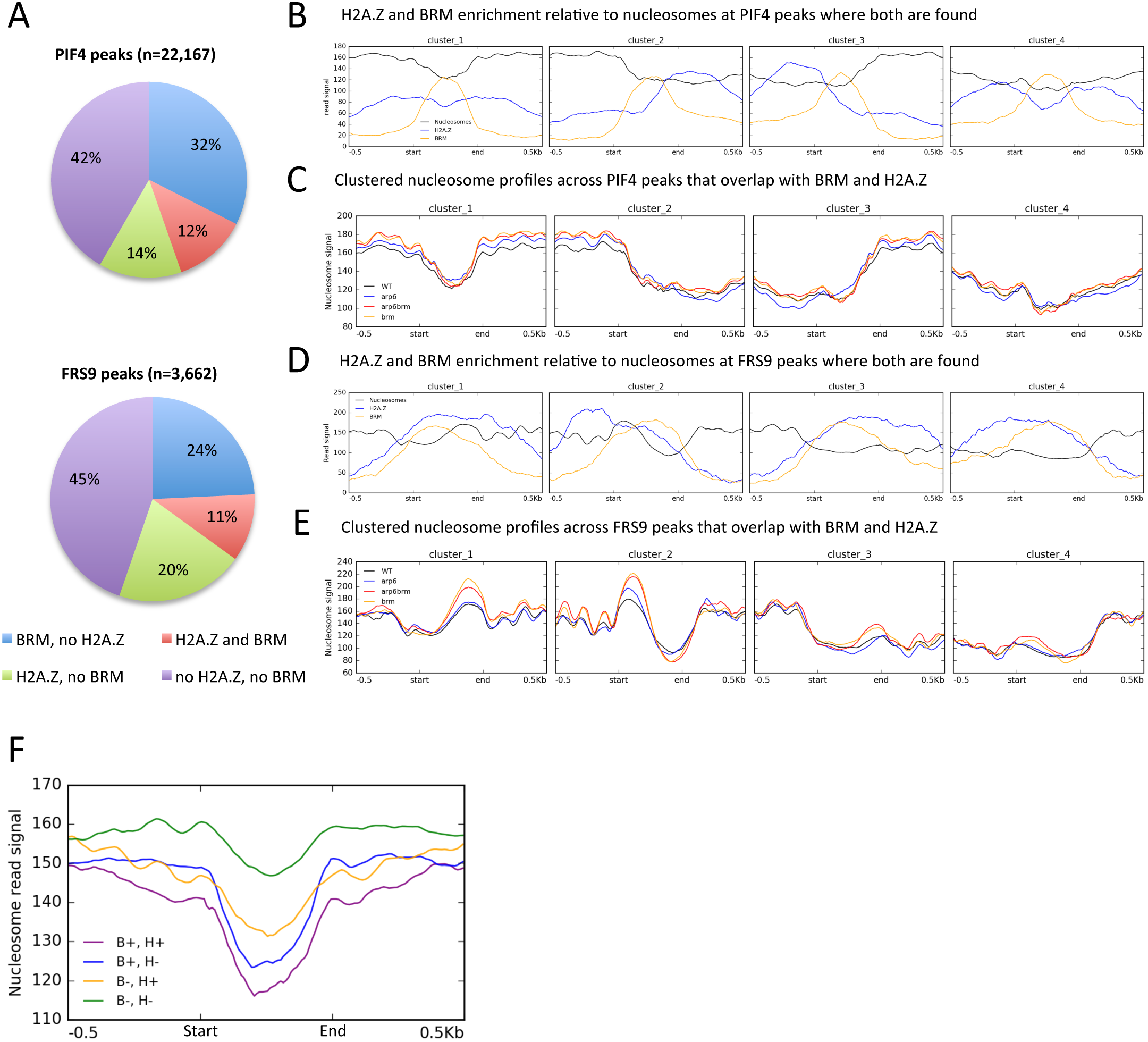
H2A.Z and BRM interact with PIF4 binding sites and contribute to nucleosome stability at FRS9 binding sites. **(A)** Pie charts display the percentage of PIF4 peaks (n=22,167, top) or FRS9 peaks (n=3,662, bottom) that overlap with BRM and not H2A.Z (blue), BRM and H2A.Z (red), H2A.Z and not BRM (green), or neither BRM nor H2A.Z (purple) ChIP-seq peaks in WT. **(B)** Average profile plots showing the BRM (orange) and H2A.Z (blue) IP signals after normalizing to input plotted along with WT nucleosome patterns (black). Signals are subdivided into 4 K-means clusters based on nucleosome patterns. **(C)** Average K-means clustered profile plots showing nucleosome reads from MNase-seq experiments across PIF4 ChIP-seq binding sites scaled to 500 bp regions. Plots show nucleosome profiles from WT (black), *arp6* (blue), *arp6;brm* (red), and *brm* (orange) plants. The same plots were performed for **(D)** BRM/H2A.Z ChIP-seq read signals and **(E)** nucleosome read signals at FRS9 binding sites. **(F)** Average profile plot showing the WT nucleosome patterns at PIF4 sites that have both BRM and H2A.Z (B+, H+; purple), BRM and not H2A.Z (B+, H-; blue), H2A.Z and not BRM (B-, H+; orange), or no BRM nor H2A.Z (B-, H-; green).

To evaluate whether chromatin organization surrounding PIF4 binding sites is dependent on H2A.Z or BRM, we plotted the nucleosome read signal from WT plants, *arp6* mutants, *brm* mutants, and *arp6;brm* double mutants across size-scaled PIF4 binding sites (Fig. 8C). We specifically plotted nucleosome read signals across PIF4 peaks that overlap with BRM ChIP-seq peaks (without H2A.Z), H2A.Z ChIP-seq peaks (without BRM), both, or neither (Fig. 8C and F, Fig. S7A-C). By K-means clustering the nucleosome profiles around PIF4 sites into four different clusters, we measured nucleosome changes at binding sites that are flanked by less accessible chromatin on one or both sides as well as sites that are found in more accessible chromatin (Fig. 8C, Fig. S7A-C). However, there are no clear differences in nucleosome occupancy around PIF4 sites when comparing the mutant nucleosomes to those in WT (Fig. 8C, Fig. S8A-C). This implies that BRM and H2A.Z do not necessarily impede or facilitate accessibility to PIF4 binding sites.

We did observe differences in the chromatin architecture inherent to the PIF4 binding sites that are associated with either BRM or H2A.Z. The clustered profiles demonstrate that more distinct nucleosome peaks flank PIF4 binding sites found in combination with H2A.Z sites (Fig. 8C and F and S7B). This suggests that well-phased nucleosomes surround PIF4 binding sites where H2A.Z is found. PIF4 binding sites found with BRM tend to be more accessible regions, consistent with BRM localizing to NDRs (Fig. 8C and S8A). We also compared WT nucleosome patterns at all PIF4 sites divided into whether they overlap with BRM alone, H2A.Z alone, both, or neither. It appears that BRM and h2a.Z localization additively correlate with more open chromatin conformations at PIF4 sites (Fig. 8F). This observation suggests that both function at more open PIF4 binding sites rather than maintaining a closed chromatin conformation. We also evaluated how the accessibility of PIF5 binding sites is affected in *arp6, brm* and *arp6;brm* mutants and discovered similar results as what we observed for PIF4 sites (Fig. S6). This is consistent with the fact that PIF4 and PIF5 have overlapping functions (De Lucas and Prat 2014). Thus, our results indicate that BRM localizes to PIF4 and PIF5 sites that are accessible, H2A.Z correlates with PIF4 and PIF5 sites within well-phased nucleosomes, and accessibility to PIF4 and PIF5 sites are not dependent on BRM and H2A.Z. Since there are no changes in the accessibility of these PIF binding sites in *brm* mutants, BRM may be recruited to these sites after PIF4 or PIF5 binds and another factor makes the region available.

### Nucleosome organization at FRS9 sites is dependent on BRM and H2A.Z

FAR1-Related Sequence 9 (FRS9), a member of the FRS far-red light responsive TF family was also predicted to interact with 6 of our classes of DE H2A.Z-BRM target genes based on our motif discovery analysis (Lin et al. 2004). Little is known about FRS9 function, but it is expressed in young rosette tissue and regulates the inhibition of hypocotyl elongation by red light (Lin et al. 2004). Also, other FRS9 paralogs bind PIF4 target genes to repress their transcription (Ritter et al., 2017). Using publically available FRS9 binding sites from DAP-seq experiments (O'malley et al., 2016), we tested whether nucleosome organization at FRS9 binding sites is dependent on H2A.Z or BRM. For this analysis, we plotted MNase-seq nucleosome signals from *arp6, brm, arp6;brm* and WT plants across sized-scaled FRS9 binding sites. Similar to how PIF4 and PIF5 were analyzed, we divided FRS9 binding sites into those sites overlapping H2A.Z ChIP-seq peaks (without BRM), overlapping BRM ChIP-seq peaks (without H2A.Z), overlapping both, or neither (Fig. 8A, Fig S8). In contrast to how BRM and H2A.Z showed more peripheral interactions at PIF4 binding sites, BRM and H2A.Z have a great degree of overlap at the FRS9 binding sites that overlap with both BRM and H2A.Z (Fig 8D). This may indicate a more coordinated function between BRM and H2A.Z in nucleosome organization at these sites.

Since relationships between nucleosomes and TF binding sites do not follow a simple presence-absence pattern, we evaluated how 4 K-means clustered nucleosome patterns associated with FRS9 binding sites are affected in *arp6, brm*, or *arp6;brm* double mutants compared to those in WT plants. At FRS9 sites overlapping with both BRM and H2A.Z peaks, there was an increase in nucleosome occupancy in the *brm* and *arp6;brm* mutants with a notable subset of nucleosomes that also have increased occupancy in *arp6* mutants (Fig. 8E and Fig. S8D-F). These occupancy changes demonstrate that both factors can contribute to nucleosome destabilization at FRS9 binding sites (Fig 8E).

These changes in occupancy were not detected at FRS9 sites that have neither H2A.Z nor BRM present (Fig. S7F), demonstrating that changes in nucleosome stability observed in the mutants can be attributed to losing H2A.Z and BRM. To better understand the individual contributions of BRM and H2A.Z, we plotted the average nucleosome signal in the *arp6, brm* and *arp6;brm* mutants and WT plants at FRS9 sites that overlapped with BRM and not H2A.Z and, conversely, those that overlap H2A.Z and not BRM. In the *brm and arp6;brm* mutants, FRS9 sites overlapping with BRM but not H2A.Z had increased occupancy for bordering nucleosomes, but a decrease in occupancy within the more accessible regions of FRS9 binding sites (Fig. S8D, clusters 2 and 3). Since we see an increase and decrease in nucleosome occupancy so close together, BRM may be responsible for moving nucleosomes from the bordering regions into the FRS9 binding sites to maintain more specific control of FRS9 or other TFs binding there. Both the peaks that are associated with H2A.Z but not BRM and those that have H2A.Z and BRM display more nucleosome phasing than what is observed in the peaks that do not have H2A.Z, similar to the nucleosome patterns we observed surrounding PIF4 binding sites (Fig. 8C and E and Fig. S8E). Since this phasing is not disrupted in the *arp6* mutants, but rather we observed an increase in the occupancy of the already well-positioned nucleosomes, H2A.Z may be needed at FSR9 sites to modulate chromatin accessibility in response to these already well-positioned nucleosomes (Fig. 8E).

Clustering FRS9 binding sites into four categories with K-means clustering allowed us to see the distribution of well-positioned nucleosomes at either side of the FRS9 binding sites (Fig. 8E). The nucleosome position that experiences the most dynamic changes at FRS9 sites in the mutant lines corresponds with nucleosomes that would normally contain H2A.Z (Fig 8D and E). H2A.Z appears to contribute to nucleosome occupancy, since we see increased nucleosome occupancy in *arp6* and *arp6;brm* mutants at FRS9 sites with H2A.Z but not BRM (Fig S7E). Although H2A.Z was not sufficient to destabilize nucleosomes in a way that resulted in nucleosome occupancy changes in the *arp6* mutants where H2A.Z overlaps with BRM, its presence could still correlate with and contribute to the role of BRM destabilizing nucleosomes. However, we see that nucleosomes at many of the borders of FRS9 peaks still accumulate in the *brm* mutants even at FRS9 binding sites where BRM localizes without H2A.Z (Fig. S7D). Thus, it seems that H2A.Z is not necessary for BRM to destabilize nucleosomes at FRS9 binding sites.

## DISCUSSION

### H2A.Z and BRM have similar and redundant roles as well as antagonistic roles in regulating transcription

Originally, we set out to test the hypothesis that BRM antagonizes the activating function of H2A.Z by stabilizing or repositioning nucleosomes which was first proposed in response to their antagonistic relationship in regulating *FLC* transcription (Farrona et al., 2011). Through the work reported here, we learned that the relationship between BRM and H2A.Z in transcriptional regulation is not a simple antagonism, but includes several different relationships. We identified gene sets where BRM antagonizes the repressive function and the activating function of H2A.Z (Classes 7 and 8), and reciprocally, where H2A.Z antagonizes the activating and repressive functions of BRM (Classes 5 and 6). We also identified genes that depend on either ARP6 or BRM to modulate transcript level (Classes 1 and 2) and genes where the additive function of both factors contributes to transcriptional repression or activation (Classes 3 and 4).

### H2A.Z levels in chromatin are independent of BRM

One hypothesis that would explain how BRM and H2A.Z coordinately or antagonistically regulate transcription is that one factor may regulate the ability of the other factor to associate with the loci that they both target. Others have observed H2A.Z protein levels increase in nuclear fractions in RNAi knock down plants for the BAF60 SWI2/SNF2 subunit, however they measured nuclear H2A.Z levels not necessarily H2A.Z levels in chromatin (Jegu et al., 2014). Since *brm* mutants do not have a consistent increase or decrease in H2A.Z-containing nucleosome levels in our ChIP-seq experiments (Fig. S1A), our results indicate that BRM does not affect H2A.Z levels in chromatin. An alternative hypothesis would be that H2A.Z and BRM interact to regulate transcription by H2A.Z recruiting BRM, because H2A.Z plays a role in recruiting the SWI2/SNF2 complex to at least one locus in human cells (Gevry et al., 2009). However, this seems unlikely since BRM and H2A.Z have a relatively small (yet significant) overlap in the genome (Fig 6A).

### BRM and H2A.Z destabilize +1 nucleosomes

At DE BRM-H2A.Z co-targeted genes, BRM localizes just upstream of the TSS and H2A.Z is enriched at the +1 nucleosome (Fig. 7B). When H2A.Z and BRM coordinately regulate gene transcription, they both contribute to +1 nucleosome destabilization (Fig. 7A, Classes 1 and 2). At other DE gene classes, BRM usually destabilizes +1 nucleosomes, while H2A.Z must contribute in other ways to transcriptional regulation. BRM and H2A.Z have been associated with +1 nucleosome stability in combination with other factors as well. Mutants for the FORGETTER1 TF that interacts with BRM perturb +1 nucleosome occupancy of genes involved in heat stress memory (Brzezinka et al., 2016). Our data showing an increase in +1 nucleosome occupancy in *brm* mutants supports a role for BRM in contributing to how FORGEtTeR1 destabilizes +1 nucleosomes after heat exposure. At the +1 nucleosomes of some heat responsive genes, H2A.Z eviction contributes to nucleosome destabilization, emphasizing a role for H2A.Z in +1 nucleosome stability (Cortijo et al., 2017). We found however that when H2A.Z is found proximal to BRM, H2A.Z tends to destabilize +1 nucleosomes of DE genes (Fig. 7A and D).

While the +1 nucleosome presents a barrier to transcription (Weber 2014), the direction of transcriptional changes observed in BRM mutants is not inherently coupled to the change in nucleosome stability caused by BRM. For example, in classes 3 and 4 of co-regulated genes, +1 nucleosome occupancy increases in BRM mutants at genes transcriptionally regulated by BRM and H2A.Z, but there are no significant transcription changes until the genome is depleted of H2A.Z-containing nucleosomes in *arp6;brm* double mutants (Fig. 7A). Changes in transcriptional regulation can also correspond with changes in the accessibility of the DNA that is associated with the +1 nucleosome without changing the nucleosome occupancy (Huebert et al., 2012). Additionally, changes in occupancy can be uncoupled from transcription changes (Mueller et al., 2017). This means that although +1 nucleosomes appear to have an increase in nucleosome occupancy in BRM mutants, and H2A.Z does not show a consistent change in nucleosome occupancy at these gene classes, the actual DNA that associates with them may have different degrees of accessibility depending on other factors such as histone modifications or interactions with other chromatin interacting proteins. In addition to contributing to transcriptional initiation, H2A.Z can help facilitate transcriptional elongation in Arabidopsis (Rudnizky et al., 2016; Weber et al 2014). The overall destabilization role of both factors in co-regulated genes may also allude to both H2A.Z and BRM contributing to transcriptional elongation rather than strictly transcriptional initiation at co-targeted genes.

### BRM destabilizes nucleosomes flanking NDRs

The specific role of BRM in chromatin regulation to date has been evaluated locus by locus, and we show here how BRM contributes to global nucleosome organization in *Arabidopsis* (Wu et al., 2015; Han 2015; Brzezinka et al., 2016). We demonstrate that BRM localizes to NDRs across the genome and is flanked by well-positioned nucleosomes whose stability often depends on BRM. These results expand on previous locus specific studies by both finding that well-positioned nucleosomes surrounding BRM binding sites is a general genomic trend and that the stability of many of these flanking nucleosomes depends on BRM (Sacharowski et al., 2015; Wu et al., 2015). Since the BAF60 subunit of the SWI2/SNF2 complex has been observed localizing to open chromatin, we provide further evidence that the SWI2/SNF2 complex binds to NDRs by finding that the BRM SWI2/SNF2 ATPase also binds to NDRs (Jegu et al., 2017). The fact that we see an increase in +1 nucleosome occupancy in the absence of BRM and H2A.Z at genes where both are needed for proper transcriptional regulation (especially Classes 1 and 2) is consistent with studies that show that both factors disrupt interactions between DNA and nucleosomes (Schnitzler et al., 2001; Rudnizky et al., 2016). It is interesting to note that although BRM localizes to NDRs, it appears that other factors establish the open confirmation of these regions and BRM may further modulate how other factors interact with the regions as we observed at PIF4 and PIF5 binding sites (Fig 8C and E).

### Other chromatin factors may contribute to the roles of BRM and H2A.Z in chromatin organization and transcriptional regulation

Some of the changes that we observed may not be due to the specific catalytic functions of BRM or inherent properties of H2A.Z incorporation into chromatin by the SWR1 complex but rather be contributed by other chromatin regulating factors that interact with them. The SWI2/SNF2 complex is known to interact with a histone acetyl transferase (HD2C), a H3K27me3 histone demethylase (REF6), and potentially the ISWI CRC (Brzezinka et al., 2016; Buszewicz et al., 2016; Li et al., 2016). BRM also antagonizes the function of the Polycomb Repressive Complex 2, so some of the nucleosomal changes we observe may not be due to a direct contribution by BRM but a result of nucleosomal changes that come with Polycomb repressive complex associated silencing activity (Li et al., 2015a). Additionally, interchanging subunits of the SWI2/SNF2 complex can confer unique functions to modulate specific developmental processes (Vercruyssen et al., 2014; Sacharowski et al., 2015). This could mean that some of the variability in BRM’s role in chromatin regulation could correspond with which SWI2/SNF2 subunits co-localize with it. BRM and the paralogous SWI2/SNF2 ATPase SPLAYED have both unique and redundant roles in *Arabidopsis*, so some contributions from BRM that are redundant with SPLAYED will be obscured from our analyses (Bezhani et al., 2007).

In other organisms, post-translational modifications to H2A.Z, such as ubiquitination and acetylation, have been shown to correlate with the role of H2A.Z in transcriptional repression and activation, respectively (Marques et al., 2010; Dalvai et al., 2012; Valdes-Mora et al., 2012). Assuming that similar post-translational modifications to H2A.Z exist in *Arabidopsis*, they likely contribute to some of the variability in nucleosome positioning and stability that we describe for H2A.Z. However, more work is still needed to create a clear and comprehensive description of how histone-modifying enzymes interact with H2A.Z and SWR1 to affect chromatin organization and regulate transcription.

Although many of the SWR1 complex subunits are shared with other CRCs, ARP6 is unique to the SWR1 complex and is essential for proper H2A.Z incorporation (Deal et al., 2007; Lu et al., 2009; Hargreaves and Crabtree 2011). This highlights the fact that the primary function reported for ARP6 in *Arabidopsis* is to incorporate H2A.Z into chromatin as part of the SWR1 complex (Deal et al., 2007; Sura et al., 2017). Thus, we used *arp6* mutants as a proxy for H2A.Z mutants in this study. ARP6 does however have functions independent of h2a.Z in yeast to localize some chromatin regions to the nuclear periphery, and some SWR1 complex subunits in *Arabidopsis* appear to have non-overlapping roles in regulating defense response (Yoshida et al., 2010; Berriri et al., 2016). Therefore, we specifically focused our analyses on regions of the genome that normally contain H2A.Z and were depleted of H2A.Z-containing nucleosomes in *arp6* mutants, thus excluding any effects from ARP6 functions that may be independent of H2A.Z and, conversely, effects from H2A.Z that may be ARP6-independent. However, it is still possible that some of our observations describe ARP6 function in addition to H2A.Z function, since we cannot parse the individual contributions of ARP6 and H2A.Z in our study.

### BRM and H2A.Z interact with binding sites for light responsive TFs

Based on our GO analyses, H2A.Z and BRM regulate transcriptional networks of genes that are involved in defense, temperature and light responses as well as growth (Fig. 2). This supports the idea that both H2A.Z and BRM contribute to the balance between normal growth and responses to stimuli. More specifically, overlapping DE target genes suggest that H2A.Z and BRM are important to integrate signals and regulate transcription in response to light stimuli. Since we show that BRM and H2A.Z co-localize with FRS9, PIF4, and PIF5 binding sites (Fig. 8 and Fig. S7), our findings indicate that interacting with light responsive TFs is one way that H2A.Z and BRM respond to light stimuli.

A relationship between FRS9 with either BRM or H2A.Z had not been observed before, but relationships between different PIF TFs and either H2A.Z or the SWI2/SNF2 complex have been reported previously. Yeast two-hybrid experiments demonstrate that BRM itself actually interacts with PIF1 and to some extent with PIF3 and PIF4 (Zhang et al. 2016), and mass spectrometry experiments show that the SWI2/SNF2 complex associates with PIF1 and 3 as well (Efroni et al., 2013). PIF1 is at least partly responsible for recruiting BRM to particular loci, however whether PIF4 and PIF5 TFs interact with BRM or H2A.Z *in vivo* was previously unknown (Zhang et al., 2017a; Efroni et al., 2013). In addition to the SWI2/SNF2 complex, H2A.Z and PIF proteins have a moderate genetic overlap in regulating flowering timing and growth in response to temperature changes, but the details of this relationship are not well understood (Wigge 2013; Galvao et al., 2015). By finding PIF TF binding sites enriched at genes co-regulated by and H2A.Z and BRM, we expand on these previously described relationships and provide resources to further explore their interactions.

Our finding of PIF TF binding at DE BRM-targeted genes provides further support for these interactions between the SWI2/SNF2 complex and PIF TFs (Table S3, Fig. 8 and Fig. S7). We also discovered that chromatin accessibility at PIF4 and PIF5 TF binding sites is not dependent on BRM nor H2A.Z (Fig. 8 and Fig. S7). Although BRM is not necessary for nucleosome organization surrounding the PIF TF binding sites, BRM may act to antagonize the function of PIF4/5 at sites where they both bind. Setting the precedent for this, the BAF60 subunit of SWI2/SNF2 competes with PIF4 for binding sites to oppose its role in hypocotyl elongation (Jegu et al., 2017).

Although *in vitro* work shows that the SWI2/SNF2 complex from other organisms repositions nucleosomes toward bound TFs to evict them (Li et al., 2015b), we did not see changes in nucleosome occupancy or positioning at PIF TF binding sites in *brm* mutants to suggest that this is the case for these TF binding sites. Alternatively, FRS9 sites can be occupied by nucleosomes and both H2A.Z and BRM regulate nucleosome occupancy at FRS9 binding sites (Fig 8D). Expanding what we know about FRS9 binding sites, we demonstrate that they overlap with BRM and H2A.Z in the genome and are found at target genes distributed across the 8 classes of DE co-targets of H2A.Z and BRM.

In other organisms, the SWI2/SNF2 CRC and H2A.Z both contribute to enhancer function to regulate transcription by regulating where TFs bind (Euskirchen et al., 2011; Brunelle et al., 2015; Alver et al., 2017). Recent work also suggests that H2A.Z may function at enhancers in *Arabidopsis* (Dai et al., 2017). However as of yet, only a small number of enhancers have been identified and characterized in plants (Zhu et al., 2015). The fact that BRM localizes to NDR more distal to TSSs and that H2A.Z and BRM are associated with TF binding sites and transcriptional regulation may indicate a role for BRM and H2A.Z in enhancer regulation in plants as well.

### Large deletions in SWR1 *mutants may account for an over estimation of nucleosome occupancy decreases*

To our knowledge no one has ever reported that there are a considerable number of large genomic deletions in SWR1 mutants in *Arabidopsis*. Our discovery of these deletions is in line with the roles of the SWR1 complex and H2A.Z in maintaining genome stability and previous reports that specifically show that *arp6* mutants have a greater crossover density, are more susceptible to DNA damage, and have meiotic defects (Choi et al., 2013; Rosa et al., 2013). Contrary to our observations of nucleosome occupancy, other groups have reported a general decrease in nucleosome occupancy in *arp6* mutants, which could, in part, be due to changes in the genome rather than changes to chromatin organization (Dai et al., 2017). In our analysis of nucleosome occupancy, we controlled for the loss of genomic regions in *arp6* mutants so that we did not erroneously report a deleted region as a decrease in nucleosome occupancy. Future studies that measure chromatin accessibility in SWR1 mutants should take care to account for similar genomic differences.

### 3-D nuclear organization in Arabidopsis may involve H2A.Z and BRM

While BRM and H2A.Z localize to a large portion of genes in the genome, only a fraction of these target genes is differentially expressed in the mutants, which is consistent with observations from previous studies of BRM (Li et al., 2016). In addition to how they directly impact the remodeling of individual nucleosomes, the roles of BRM and H2A.Z in transcriptional regulation may contribute to or be a consequence of larger nuclear organization of chromatin. Transcription can be oversimplified if viewed as an isolated linear process of initiation, elongation, and termination proceeding down a DNA molecule. In reality, transcription is one of many processes that take place in a highly regulated chromatin environment that is organized in an intricate 3-dimensional space within the nucleus (Vergara and Gutierrez 2017; Barneche and Baroux 2017). Both H2A.Z and the SWI2/SNF2 complex have been implicated in regulating larger scale nuclear organization in other organisms, contributing to chromatin looping and chromosome localization within the nucleus (Yoshida et al., 2010; Light et al., 2010; Maruyama et al., 2012; Kitamura et al., 2015; Imbalzano et al., 2013). The fact that H2A.Z and the SWR1 complex associate with nuclear scaffold/matrix attachment regions in Arabidopsis suggests that similar functions for both are yet to be described in plants (Lee and Seo 2017). Likewise, the SWI2/SNF2 complex has been implicated in chromatin looping in *Arabidopsis* and other organisms, as well as *in vitro* (Jegu et al., 2014, Kim et al., 2009, Bazett-Jones et al., 1999). Therefore, changes in higher order structure may be affected by depleting the plants of BRM and H2A.Z-containing nucleosomes, which could explain some of the variable changes in nucleosome stability we observed in *brm* and *arp6* mutants and why some H2A.Z and BRM associated nucleosomes do not dramatically change. Additionally, plants respond to changes in light signals with chromatin de-condensation and nuclear reorganization, so a deeper mechanistic understanding of BRM and/or H2A.Z may elucidate how chromatin changes occur on a larger scale in response to changes in light or other stimuli (Van Zanten et al., 2010; Bourbousse et al., 2015).

#### Concluding statements

Within the nucleus, combinatorial effects from a range of factors regulate chromatin organization in different contexts. This can make it difficult to understand the extent to which any one factor contributes to chromatin organization as a whole. *In vitro* studies work to simplify the system to understand individual chromatin-influencing components, but they are far removed from the constant flux of regulatory pressures that a locus experiences *in vivo*. In our study, we attempted to parse the chromatin regulatory contributions of H2A.Z and BRM *in vivo* and chose to simplify our approach by identifying and then specifically evaluating direct target loci of H2A.Z and BRM where they antagonistically or coordinately regulate transcription through multiple regulatory relationships (Fig 1E). We found that not only do H2A.Z and BRM work at co-targeted genes to positively and negatively regulate genes, but some of their roles are functionally redundant. In addition, we identified genes where H2A.Z and BRM act either negatively or positively to affect transcript level in ways that are opposed by the other factor. We discovered that BRM contributes more to stabilizing nucleosomal changes where it directly binds to chromatin with more destabilizing effects on flanking nucleosomes. However, H2A.Z-containing nucleosomes have no clear enrichment for a specific type of nucleosome dynamic when it is found on its own or in association with BRM. At co-targeted DE genes, BRM and H2A.Z contribute to nucleosome stability to varying degrees, but they appear to both regulate +1 nucleosome occupancy where either is required for transcriptional regulation (Fig. 7). Some of the variability in H2A.Z and BRM function may be explained by their interactions with specific TFs, such as the three TFs we identified (Fig 8). While these datasets help us better survey how both H2A.Z and BRM contribute to transcription and nucleosome organization, more cell-type and locus-specific studies are needed to understand their full contribution to chromatin level regulation.

These genetic dissections indicate that the relationship between BRM and H2A.Z is more complicated than one property of each factor contributing to or antagonizing a single function of the other. Therefore, the influence of additional factors must make the roles of H2A.Z and BRM necessary to regulate transcription levels in different contexts. Further *in vivo* genetic and molecular studies will help us identify which factors define the context dependent functions of BRM and H2A.Z, while *in vitro* studies would help simplify the experimental system and define the direct interactions between H2A.Z and BRM as well as additional identified factors. This highlights the challenge we face in chromatin research to create simple enough systems to understand the true complexity of how individual chromatin associating factors function on a sophisticated chromatin template within the nucleus (Probst and Mittelsten Scheid 2015).

## MATERIALS AND METHODS

### Plant material

We used previously characterized *Arabidopsis* T-DNA insertion lines *arp6–1* (GARLlC_599_G03; Deal et al., 2005) and *brm-1* (SALK_030046, Hurtado et al., 2006) and genotyped the strains using primers described previously (Deal et al., 2005; Hurtado et al., 2006). The *arp6–1;brm-1* mutant was generated from genetic crosses of *arp6–1* homozygous and *brm-1* heterozygous lines. Plants were sown on soil, stratified at 4 °C for two days, and then moved to grow at 20 °C in long day light conditions (16 hr light/8 hr dark). Above ground plant tissue for all genomic experiments was collected at 10 hrs after dawn from 4–5 leaf developmentally staged plants (Boyes et al., 2001) from the following genetic backgrounds: WT (collected 12–13 days post stratification (dps)); *arp6–1* (12–14 dps); *brm-1* (13–16 dps); and *arp6;brm* (16–24 dps, delayed collection due to delayed germination). One cotyledon was removed from each plant to use for genotyping with the Phire^TM^ Plant Direct PCR Kit (Thermo Scientific).

### RNA-seq material

Three plants each for three biological replicates of 4–5 leaf developmentally staged above soil seedling material were collected and pooled for WT, *arp6, brm*, and *arp6;brm* plants. RNA was isolated using the Spectrum Plant Total RNA Kit (Sigma) and incubated at 37 °C for 30 min with DNase to remove DNA using the Turbo DNA-free kit (Ambion). The integrity of the RNA was confirmed on a 2% agarose gel in 1x TAE visualized with GELRED nucleic acid stain (Sigma), and the samples were quantified with a spectrophotometer. Libraries were prepared from 100 ng of RNA from each sample using the Ovation RNA-seq for Model Organisms kit (NuGEN), which is a strand specific library preparation kit that depletes the transcripts of rRNA. Libraries were quantified with qPCR (NEB), pooled, and sequenced with the Illumina NextSeq500 to generate paired-end 36-nt sequence reads.

### RNA-seq data analysis

Sequencing reads were mapped to the TAIR10 *Arabidopsis thaliana* reference genome using Tophat2 (using the second strand option and default parameters), generating an average of 75.5M mapped reads per library. The accepted hits file was name-sorted (option –n) rather than position sorted and indexed using SAMtools (Trapnell et al., 2012; Li et al., 2009). Read counts were quantified for each exon using the htseq-counts program, with name order and strict intersection options (Anders et al., 2015). Differential expression was calculated using edgeR software (Robinson et al., 2010; Mccarthy et al., 2012). Differentially expressed genes were determined with a false discovery rate (FDR) cutoff of <0.2 and a log_2_ fold change of ± 0.6 (~1.5 x fold change). GO terms were generated using AgriGO for the total set of genes that were DE in the mutants relative to WT plants. GeneCodis was used to analyze GO terms for DE direct target genes of H2A.Z and BRM (Carmona-Saez et al., 2007; Nogales-Cadenas et al., 2009; Tabas-Madrid et al., 2012; Tian et al., 2017). These two separate programs were used for GO analyses based on how generally (AgriGO) or specifically (GeneCodis) they summarized the overlap between gene lists.

### ChIP-seq material

For ChIP-seq experiments, we collected at least 0.5 g of tissue from two biological replicates each of WT, *arp6–1*, and *brm-1* plants. (WT, 12–13 dps; *arp6*, 12–14 dps; *brm*, 13–16 dps; *arp6;brm* 16–24 dps). Above ground developmentally staged 4–5 leaf plant tissue was collected at 10 hrs after dawn, cross-linked as described previously (Gendrel et al., 2005), frozen, and ground in liquid nitrogen. Nuclei were isolated as previously described (Gendrel et al., 2005). Chromatin was sonicated using a Bioruptor^®^ (Diagenode) (40 min on high (45 sec on/ 15 sec off)). Each sample was diluted in 1.1 mL of ChIP dilution buffer (described in Gendrel et al., 2005) and 50 μl was saved as the input sample. Then H2A.Z-containing chromatin was immunoprecipitated from the 1.1 mL of chromatin solution using 2 μg of H2A.Z antibody purified to specifically recognize unmodified H2A.Z peptides (Deal et al., 2007). The chromatin solution was incubated with the H2A.Z antibody for 2 hr then for 1 more hour in combination with 60 μl of Dynabeads™ Protein-A magnetic beads (Invitrogen). DNA collected from the immunoprecipitation and from the inputs was purified using 1.8x volume of SPRI beads (Beckman Coulter) then quantified with Quant-IT^TM^ Picogreen^®^ dsDNA Assay Kit (Invitrogen). Sequencing libraries were prepared from 1 ng of DNA per sample with the Accel-NGS^®^ 2S Plus DNA Library kit (Swift Biosciences) and sequenced on an Illumina NextSeq500 using 76 nt single-end reads.

### ChIP-seq data analysis

ChIP-seq reads were mapped to the TAIR10 *A. thaliana* reference genome with Bowtie2, using default parameters (Langmead and Salzberg 2012). An average of 13.9 M reads were converted to binary files, sorted, indexed and quality filtered (with the -q 2 option) using SAMtools software (Li et al., 2009). H2A.Z peaks were called with Homer findpeaks software, using options “style histone” and “-region” (Heinz et al., 2010). H2A.Z peaks from two biological replicates were intersected to find the regions that were called in both replicates for each genotype using Bedtools software (Quinlan 2014). H2A.Z peaks in WT that overlapped with H2A.Z peaks called in *arp6* mutants with less than a 2-fold difference in enrichment between the two genotypes were removed from the analysis to ensure that the datasets analyzed represent ARP6-dependent H2A.Z peaks. These were the H2A.Z peaks we used throughout the study. We integrated BRM-GFP ChIP-seq peaks into our analysis from a previously published data set (Li et al., 2016). Also, we used previously published ChIP-seq data for PIF4 (AT2G43010) (SRX1005830) and PIF5 (AT3G59060) (SRX1495297) (Pedmale et al., 2016) and DAP-seq peaks for FRS9 (O'malley et al., 2016). H2A.Z and BRM ChIP-seq peaks were annotated based on the genes that they overlapped (-u ODS option) or were assigned to the nearest TSS (-u TSS option) using PeakAnnotator software (Salmon-Divon et al., 2010). Before preparing bigwig files, we first used the SAMtools view command (with option –s) to scale data sets so that all samples had same number of reads. We also combined the two biological replicates with SAMtools merge. Using the deepTools software suite, we then prepared bigwig files using default parameters for the bamCoverage program, then we subtracted the input signal from the ChIP signals by 10 bp bins with the bamCompare command for each genotype (Li et al., 2009; Li 2011; Ramirez et al., 2014). Heatmaps and average profile plots were generated from these bigwigs using deepTools computeMatrix, plotHeatmap, and plotProfile programs (Ramirez et al., 2014).

### MNase-seq material

Tissue from two biological replicates of 100 mg of pooled above ground 4–5 leaf stage plants was collected from WT, *arp6–1, brm-1, and arp6–1;brm-1* plants grown on soil in long day light conditions (16 hr light/ 8 hr dark). Nuclei were isolated as described previously (Gendrel et al., 2005). After purification, nuclei were resuspended in 500 μl of TM2 solution (10 mM Tris (pH 7.5), 2 mM MgCl_2_, and 1X Roche Complete protease inhibitor tablet). We spun nuclei down at 3,000 × g for 10 min then removed the supernatant and re-suspended the pellet in 500 μl of MNase reaction buffer (16 mM Tris-Cl (pH 8.0), 50 mM NaCl, 2.5 mM CaCl_2_, 1 mM EDTA, Protease inhibitor tablet). Samples consisting of 500 μl of nuclei were incubated with 7.5 U MNase for 7.5 min at 37 °C, and then the reaction was stopped by adding EDTA to a final concentration of 10 mM. Nuclei were lysed by adding sDS (to 1% of the final sample volume). The solution was mixed and spun down at 1,300 x g for 3 min to remove insoluble debris. After moving the supernatant to a new tube, samples were treated with RNase A (1 mg/mL, Ambion) and then with Proteinase K (Invitrogen) to remove RNA and proteins, respectively. DNA fragments were purified with MinElute PCR purification kit (Qiagen). To purify nucleosome associated DNA fragments that were <400 bp, we used a 0.6x bead-to-sample ratio of SPRI beads prepared as described previously (Faircloth and Glenn 2014). Sequencing libraries were prepared with the ThruPLEX^®^ DNA-Seq Kit (Rubicon Genomics), using 25 ng of MNase-digested DNA as the input. Purified, indexed libraries were pooled and sequenced on an Illumina NextSeq500 generating 76 bp paired-end reads.

### MNase-seq data analysis

Sequence reads were mapped to the TAIR10 *Arabidopsis thaliana* reference genome using Bowtie2 (using default parameters except for –p 6) and were then further sorted and indexed using SAMtools (Li et al., 2009; Langmead and Salzberg 2012). We filtered reads using the SAMtools view command, with the –q 2 option to filter for quality and option –f 0×02 to filter for properly paired reads. Libraries were subsampled using the SAMtools –s parameter to normalize all samples to the same number of reads (30.46 M reads). We analyzed the mapped reads from each biological replicate for each of the four genotypes to generate nucleosome peak files and nucleosome occupancy wiggle files using the DANPOS2 dpos program (Chen et al., 2013). Values in the *.allPeaks.xls file output from the dpos program were used to determine dynamic nucleosomes. These dynamic nucleosomes are defined as those with a FDR <0.05 for the difference between the occupancy value at the summit position of a point of difference in control and treatment samples (point_diff_FDR <0.05) and then individual types of dynamic nucleosome changes were described with the following additional criteria. Fuzziness scores were defined as the standard deviation of read positions in each peak. Significant nucleosome fuzziness changes were defined as those with a FDR of <0.05 for the difference between WT and mutant fuzziness scores (fuzziness_diff_FDR <0.05). Significant occupancy changes were defined as those with a FDR of <0.05 of the difference between the occupancy value at the peak summit position in the WT and the mutant (smt_diff_FDR <0.05). Position shifts were defined as a 20–95 bp difference in peak summit position between WT and mutant nucleosomes (treat2control_dis 20–95 bp). To measure nucleosome occupancy across the genome, we converted the DANPOS generated wiggle files to bigwig files using the wigToBigWig software (UCSC). Heatmaps were generated from these bigwig files using deepTools software: computeMatrix, plotHeatmap, and plotProfile programs (Ramirez et al., 2014).

### Identifying deletions in *arp6* mutants

We isolated genomic DNA from mature rosette leaf material (~5 mg per plant) from 50 pooled *arp6* mutants using a standard phenol:chloroform extraction followed by ethanol precipitation. We used 1 μg of sonicated DNA to prepare a sequencing library (NEXTflex Rapid DNA-seq (option 2), Bioo Scientific), and then sequenced on an Illumina HiSeq2000, generating 125 bp paired-end reads. Sequenced reads were mapped using BWA mem software, indexed and quality sorted (-s option) using SAMtools, and randomly subsetted using a python script, leaving 61.5 M mapped reads (Li and Durbin 2009; Li et al., 2009; Li 2011). Deletions were called in the *arp6* mutant using CNVnator (v0.3.3) software using bin sizes of 100 bp (Abyzov et al., 2011).

### Motif Enrichment Analysis

ATAC-seq transposase hypersensitivity sites (THSs) from mesophyll cells were identified and annotated previously (Sijacic et al., 2017). We used python scripts to pull out the annotated mesophyll THSs that were associated with genes from our 8 differentially expressed BRM and H2A.Z target gene classes (defined in Fig.1 D). We scaled all THSs to be 150 bp in width and then used the Regulatory Sequence Analysis Tool for plants (RSAT plants) to obtain the corresponding DNA sequences and mask any repeated sequences (Medina-Rivera et al., 2015). Motifs that were enriched in our lists of THSs were discovered using DREME and MEME programs from the MEME-ChIP suite then paired with TFs predicted to bind to the motifs using the Tomtom program (Bailey et al., 2009) THS sequences were compared to both the CIS-BP and DAP-seq TF binding databases for these analyses (Weirauch et al., 2014; O'malley et al., 2016). Motifs and their associated TF were considered significant they had an E-value of < 0.05.

### Accession Numbers

All high throughput sequencing data described in this paper has been deposited to the NCBI GEO database under record number (GSE108450). The BRM chIP-seq data used in our analysis is available under GEO number SRX1184288. The PIF4 and PIF ChIP-seq data used in our analyses are available under GEO numbers SRX1005830 (PIF4) and SRX1495297 (PIF5). ATAC-seq data from mesophyll cells can be found under GEO accession number GSE101940.

## Literature Cited

Abyzov A, Urban AE, Snyder M, Gerstein M (2011) CNVnator: an approach to discover, genotype, and characterize typical and atypical CNVs from family and population genome sequencing. Genome Res 21: 974–84

Alazem M, Lin NS (2015) Roles of plant hormones in the regulation of host-virus interactions. Mol Plant Pathol 16: 529–40

Allan J, Fraser RM, Owen-Hughes T, Keszenman-Pereyra D (2012) Micrococcal nuclease does not substantially bias nucleosome mapping. J Mol Biol 417: 152–64

Anders S, Pyl PT, Huber W (2015) HTSeq--a Python framework to work with high-throughput sequencing data. Bioinformatics 31: 166–9

Archacki R, Buszewicz D, Sarnowski TJ, Sarnowska E, Rolicka AT, Tohge T, Fernie AR, Jikumaru Y, Kotlinski M, Iwanicka-Nowicka R, Kalisiak K, Patryn J, Halibart-Puzio J, Kamiya Y, Davis SJ, Koblowska MK, Jerzmanowski A (2013) BRAHMA ATPase of the SWI/SNF Chromatin Remodeling Complex Acts as a Positive Regulator of Gibberellin-Mediated Responses in Arabidopsis. PLoS One 8: e58588

Archacki R, Yatusevich R, Buszewicz D, Krzyczmonik K, Patryn J, Iwanicka-Nowicka R, Biecek P, Wilczynski B, Koblowska M, Jerzmanowski A, Swiezewski S (2016) Arabidopsis SWI/SNF chromatin remodeling complex binds both promoters and terminators to regulate gene expression. Nucleic Acids Res

Bailey TL, Boden M, Buske FA, Frith M, Grant CE, Clementi L, Ren J, Li WW, Noble WS (2009) MEME SUITE: tools for motif discovery and searching. Nucleic Acids Res 37: W202–8

Barah P, B NM, Jayavelu ND, Sowdhamini R, Shameer K, Bones AM (2016) Transcriptional regulatory networks in Arabidopsis thaliana during single and combined stresses. Nucleic Acids Res 44: 3147–64

Barneche F, Baroux C (2017) Unreeling the chromatin thread: a genomic perspective on organization around the periphery of the Arabidopsis nucleus. Genome Biol 18: 97

Bazett-Jones DP, Cote J, Landel CC, Peterson CL, Workman JL (1999) The SWI/SNF complex creates loop domains in DNA and polynucleosome arrays and can disrupt DNA-histone contacts within these domains. Mol Cell Biol 19: 1470–8

Berriri S, Gangappa SN, Kumar SV (2016) SWR1 Chromatin-Remodeling Complex Subunits and H2A.Z Have Non-overlapping Functions in Immunity and Gene Regulation in Arabidopsis. Mol Plant 9: 1051–65

Bevan M, Walsh S (2005) The Arabidopsis genome: a foundation for plant research. Genome Res 15: 1632–42

Bezhani S, Winter C, Hershman S, Wagner JD, Kennedy JF, Kwon CS, Pfluger J, Su Y, Wagner D (2007) Unique, shared, and redundant roles for the Arabidopsis SWI/SNF chromatin remodeling ATPases BRAHMA and SPLAYED. Plant Cell 19: 403–16

Bonisch C, Hake SB (2012) Histone H2A variants in nucleosomes and chromatin: more or less stable? Nucleic Acids Res 40: 10719–41

Bourbousse C, Mestiri I, Zabulon G, Bourge M, Formiggini F, Koini MA, Brown SC, Fransz P, Bowler C, Barneche F (2015) Light signaling controls nuclear architecture reorganization during seedling establishment. Proc Natl Acad Sci U S A 112: E2836–44

Boyes DC, Zayed AM, Ascenzi R, McCaskill AJ, Hoffman NE, Davis KR, Gorlach J (2001) Growth stage-based phenotypic analysis of Arabidopsis: a model for high throughput functional genomics in plants. Plant Cell 13: 1499–510

Brzezinka K, Altmann S, Czesnick H, Nicolas P, Gorka M, Benke E, Kabelitz T, Jahne F, Graf A, Kappel C, Baurle I (2016) Arabidopsis FORGETTER1 mediates stress-induced chromatin memory through nucleosome remodeling. Elife 5:

Buszewicz D, Archacki R, Palusinski A, Kotlinski M, Fogtman A, Iwanicka-Nowicka R, Sosnowska K, Kucinski J, Pupel P, Oledzki J, Dadlez M, Misicka A, Jerzmanowski A, Koblowska MK (2016) HD2C histone deacetylase and a SWI/SNF chromatin remodeling complex interact and both are involved in mediating the heat stress response in Arabidopsis. Plant Cell Environ

Carmona-Saez P, Chagoyen M, Tirado F, Carazo JM, Pascual-Montano A (2007) GENECODIS: a web-based tool for finding significant concurrent annotations in gene lists. Genome Biol 8: R3

Catala R, Medina J, Salinas J (2011) Integration of low temperature and light signaling during cold acclimation response in Arabidopsis. Proc Natl Acad Sci U S A 108: 16475–80

Chen K, Xi Y, Pan X, Li Z, Kaestner K, Tyler J, Dent S, He X, Li W (2013) DANPOS: dynamic analysis of nucleosome position and occupancy by sequencing. Genome Res 23: 341–51

Choi K, Zhao X, Kelly KA, Venn O, Higgins JD, Yelina NE, Hardcastle TJ, Ziolkowski PA, Copenhaver GP, Franklin FC, McVean G, Henderson IR (2013) Arabidopsis meiotic crossover hot spots overlap with H2A.Z nucleosomes at gene promoters. Nat Genet 45: 1327–36

Clapier CR, Iwasa J, Cairns BR, Peterson CL (2017) Mechanisms of action and regulation of ATP-dependent chromatin-remodelling complexes. Nat Rev Mol Cell Biol 18: 407–422

Coleman-Derr D, Zilberman D (2012) Deposition of histone variant H2A.Z within gene bodies regulates responsive genes. PLoS Genet 8: e1002988

Cortijo S, Charoensawan V, Brestovitsky A, Buning R, Ravarani C, Rhodes D, van Noort J, Jaeger KE, Wigge PA (2017) Transcriptional regulation of the ambient temperature response by H2A.Z-nucleosomes and HSF1 transcription factors in Arabidopsis. Mol Plant

Dai X, Bai Y, Zhao L, Dou X, Liu Y, Wang L, Li Y, Li W, Hui Y, Huang X, Wang Z, Qin Y (2017) H2A.Z Represses Gene Expression by Modulating Promoter Nucleosome Structure and Enhancer Histone Modifications in Arabidopsis. Mol Plant 10: 1274–1292

Dalvai M, Bellucci L, Fleury L, Lavigne AC, Moutahir F, Bystricky K (2012) H2A.Z-dependent crosstalk between enhancer and promoter regulates Cyclin D1 expression. Oncogene

de Lucas M, Prat S (2014) PIFs get BRright: PHYTOCHROME INTERACTING FACTORs as integrators of light and hormonal signals. New Phytol 202: 1126–41

Deal R, Topp CN, McKinney EC, Meagher RB (2007) Repression of Flowering in Arabidopsis Requires Activation of FLOWERING LOCUS C Expression by the Histone Variant H2A.Z. The Plant Cell Online 19: 74–83

Deal RB, Kandasamy MK, McKinney EC, Meagher RB (2005) The nuclear actin-related protein ARP6 is a pleiotropic developmental regulator required for the maintenance of FLOWERING LOCUS C expression and repression of flowering in Arabidopsis. Plant Cell 17: 2633–46

Efroni I, Han SK, Kim HJ, Wu MF, Steiner E, Birnbaum KD, Hong JC, Eshed Y, Wagner D (2013) Regulation of leaf maturation by chromatin-mediated modulation of cytokinin responses. Dev Cell 24: 438–45

Euskirchen GM, Auerbach RK, Davidov E, Gianoulis TA, Zhong G, Rozowsky J, Bhardwaj N, Gerstein MB, Snyder M (2011) Diverse roles and interactions of the SWI/SNF chromatin remodeling complex revealed using global approaches. PLoS Genet 7: e1002008

Faircloth BC, Glenn TC (2014) Protocol: Preparation of an AMPure XP substitute (AKA Serapure).

Farrona S, Hurtado L, March-Diaz R, Schmitz RJ, Florencio FJ, Turck F, Amasino RM, Reyes JC (2011) Brahma is required for proper expression of the floral repressor FLC in Arabidopsis. PLoS One 6: e17997

Galvao VC, Collani S, Horrer D, Schmid M (2015) Gibberellic acid signaling is required for ambient temperature-mediated induction of flowering in Arabidopsis thaliana. Plant J 84: 949–62

Gendrel AV, Lippman Z, Martienssen R, Colot V (2005) Profiling histone modification patterns in plants using genomic tiling microarrays. Nat Methods 2: 213–8

Gevry N, Hardy S, Jacques PE, Laflamme L, Svotelis A, Robert F, Gaudreau L (2009) Histone H2A.Z is essential for estrogen receptor signaling. Genes Dev 23: 1522–33

Han SK, Sang Y, Rodrigues A, Biol F, Wu MF, Rodriguez PL, Wagner D (2012) The SWI2/SNF2 Chromatin Remodeling ATPase BRAHMA Represses Abscisic Acid Responses in the Absence of the Stress Stimulus in Arabidopsis. Plant Cell 24: 4892–906

Hargreaves DC, Crabtree GR (2011) ATP-dependent chromatin remodeling: genetics, genomics and mechanisms. Cell Res 21: 396–420

Heinz S, Benner C, Spann N, Bertolino E, Lin YC, Laslo P, Cheng JX, Murre C, Singh H, Glass CK (2010) Simple combinations of lineage-determining transcription factors prime cis-regulatory elements required for macrophage and B cell identities. Mol Cell 38: 576–89

Hiruma K, Onozawa-Komori M, Takahashi F, Asakura M, Bednarek P, Okuno T, Schulze-Lefert P, Takano Y (2010) Entry mode-dependent function of an indole glucosinolate pathway in Arabidopsis for nonhost resistance against anthracnose pathogens. Plant Cell 22: 2429–43

Huebert DJ, Kuan PF, Keles S, Gasch AP (2012) Dynamic changes in nucleosome occupancy are not predictive of gene expression dynamics but are linked to transcription and chromatin regulators. Mol Cell Biol 32: 1645–53

Hurtado L, Farrona S, Reyes JC (2006) The putative SWI/SNF complex subunit BRAHMA activates flower homeotic genes in Arabidopsis thaliana. Plant Mol Biol 62: 291–304

Imbalzano AN, Imbalzano KM, Nickerson JA (2013) BRG1, a SWI/SNF chromatin remodeling enzyme ATPase, is required for maintenance of nuclear shape and integrity. Commun Integr Biol 6: e25153

Jegu T, Latrasse D, Delarue M, Hirt H, Domenichini S, Ariel F, Crespi M, Bergounioux C, Raynaud C, Benhamed M (2014) The BAF60 Subunit of the SwI/SNF Chromatin-Remodeling Complex Directly Controls the Formation of a Gene Loop at FLOWERING LOCUS C in Arabidopsis. Plant Cell

Jegu T, Veluchamy A, Ramirez-Prado JS, Rizzi-Paillet C, Perez M, Lhomme A, Latrasse D, Coleno E, Vicaire S, Legras S, Jost B, Rougee M, Barneche F, Bergounioux C, Crespi M, Mahfouz MM, Hirt H, Raynaud C, Benhamed M (2017) The Arabidopsis SWI/SNF protein BAF60 mediates seedling growth control by modulating DNA accessibility. Genome Biol 18: 114

John S, Sabo PJ, Johnson TA, Sung MH, Biddie SC, Lightman SL, Voss TC, Davis SR, Meltzer PS, Stamatoyannopoulos JA, Hager GL (2008) Interaction of the glucocorticoid receptor with the chromatin landscape. Mol Cell 29: 611–24

Kim SI, Bresnick EH, Bultman SJ (2009) BRG1 directly regulates nucleosome structure and chromatin looping of the alpha globin locus to activate transcription. Nucleic Acids Res 37: 6019–27

Kitamura H, Matsumori H, Kalendova A, Hozak P, Goldberg IG, Nakao M, Saitoh N, Harata M (2015) The actin family protein ARP6 contributes to the structure and the function of the nucleolus. Biochem Biophys Res Commun 464: 554–60

Lai WKM, Pugh BF (2017) Understanding nucleosome dynamics and their links to gene expression and DNA replication. Nat Rev Mol Cell Biol 18: 548–562

Langmead B, Salzberg SL (2012) Fast gapped-read alignment with Bowtie 2. Nat Methods 9: 357–9

Lee K, Seo PJ (2017) Coordination of matrix attachment and ATP-dependent chromatin remodeling regulate auxin biosynthesis and Arabidopsis hypocotyl elongation. PLoS One 12: e0181804

Lee N, Choi G (2017) Phytochrome-interacting factor from Arabidopsis to liverwort. Curr Opin Plant Biol 35: 54–60

Li C, Chen C, Gao L, Yang S, Nguyen V, Shi X, Siminovitch K, Kohalmi SE, Huang S, Wu K, Chen X, Cui Y (2015a) The Arabidopsis SWI2/SNF2 chromatin Remodeler BRAHMA regulates polycomb function during vegetative development and directly activates the flowering repressor gene SVP. PLoS Genet 11: e1004944

Li C, Gu L, Gao L, Chen C, Wei CQ, Qiu Q, Chien CW, Wang S, Jiang L, Ai LF, Chen CY, Yang S, Nguyen V, Qi Y, Snyder MP, Burlingame AL, Kohalmi SE, Huang S, Cao X, Wang ZY, Wu K, Chen X, Cui Y (2016) Concerted genomic targeting of H3K27 demethylase REF6 and chromatin-remodeling ATPase BRM in Arabidopsis. Nat Genet 48: 687–93

Li H, Durbin R (2009) Fast and accurate short read alignment with Burrows-Wheeler transform. Bioinformatics 25: 1754–60

Li H, Handsaker B, Wysoker A, Fennell T, Ruan J, Homer N, Marth G, Abecasis G, Durbin R, Genome Project Data Processing S (2009) The Sequence Alignment/Map format and SAMtools. Bioinformatics 25: 2078–9

Li H (2011) A statistical framework for SNP calling, mutation discovery, association mapping and population genetical parameter estimation from sequencing data. Bioinformatics 27: 2987–93

Li M, Hada A, Sen P, Olufemi L, Hall MA, Smith BY, Forth S, McKnight JN, Patel A, Bowman GD, Bartholomew B, Wang MD (2015b) Dynamic regulation of transcription factors by nucleosome remodeling. Elife 4:

Li Z, Gadue P, Chen K, Jiao Y, Tuteja G, Schug J, Li W, Kaestner KH (2012) Foxa2 and H2A.Z mediate nucleosome depletion during embryonic stem cell differentiation. Cell 151: 1608–16

Light WH, Brickner DG, Brand VR, Brickner JH (2010) Interaction of a DNA zip code with the nuclear pore complex promotes H2A.Z incorporation and INO1 transcriptional memory. Mol Cell 40: 112–25

Lu PY, Levesque N, Kobor MS (2009) NuA4 and SWR1-C: two chromatin-modifying complexes with overlapping functions and components. Biochem Cell Biol 87: 799–815

Luger K, Mader AW, Richmond RK, Sargent DF, Richmond TJ (1997) Crystal structure of the nucleosome core particle at 2.8 A resolution. Nature 389: 251–60

March-Diaz R, Garcia-Dominguez M, Lozano-Juste J, Leon J, Florencio FJ, Reyes JC (2008) Histone H2A.Z and homologues of components of the SWR1 complex are required to control immunity in Arabidopsis. Plant J 53: 475–87

Marques M, Laflamme L, Gervais AL, Gaudreau L (2010) Reconciling the positive and negative roles of histone H2A.Z in gene transcription. Epigenetics 5: 267–72

Maruyama EO, Hori T, Tanabe H, Kitamura H, Matsuda R, Tone S, Hozak P, Habermann FA, von Hase J, Cremer C, Fukagawa T, Harata M (2012) The actin family member Arp6 and the histone variant H2A.Z are required for spatial positioning of chromatin in chicken cell nuclei. J Cell Sci 125: 3739–43

McCarthy DJ, Chen Y, Smyth GK (2012) Differential expression analysis of multifactor RNA-Seq experiments with respect to biological variation. Nucleic Acids Res 40: 4288–97

Medina-Rivera A, Defrance M, Sand O, Herrmann C, Castro-Mondragon JA, Delerce J, Jaeger S, Blanchet C, Vincens P, Caron C, Staines DM, Contreras-Moreira B, Artufel M, Charbonnier-Khamvongsa L, Hernandez C, Thieffry D, Thomas-Chollier M, van Helden J (2015) RSAT 2015: Regulatory Sequence Analysis Tools. Nucleic Acids Res 43: W50–6

Mueller B, Mieczkowski J, Kundu S, Wang P, Sadreyev R, Tolstorukov MY, Kingston RE (2017) Widespread changes in nucleosome accessibility without changes in nucleosome occupancy during a rapid transcriptional induction. Genes Dev 31: 451–462

Narlikar GJ, Sundaramoorthy R, Owen-Hughes T (2013) Mechanisms and functions of ATP-dependent chromatin-remodeling enzymes. Cell 154: 490–503

Nogales-Cadenas R, Carmona-Saez P, Vazquez M, Vicente C, Yang X, Tirado F, Carazo JM, Pascual-Montano A (2009) GeneCodis: interpreting gene lists through enrichment analysis and integration of diverse biological information. Nucleic Acids Res 37: W317–22

O'Malley RC, Huang SC, Song L, Lewsey MG, Bartlett A, Nery JR, Galli M, Gallavotti A, Ecker JR (2016) Cistrome and Epicistrome Features Shape the Regulatory DNA Landscape. Cell 166: 1598

Pedmale UV, Huang SC, Zander M, Cole BJ, Hetzel J, Ljung K, Reis PAB, Sridevi P, Nito K, Nery JR, Ecker JR, Chory J (2016) Cryptochromes Interact Directly with PIFs to Control Plant Growth in Limiting Blue Light. Cell 164: 233–245

Probst AV, Mittelsten Scheid O (2015) Stress-induced structural changes in plant chromatin. Curr Opin Plant Biol 27: 8–16

Quinlan AR (2014) BEDTools: The Swiss-Army Tool for Genome Feature Analysis. Curr Protoc Bioinformatics 47: 11 12 1–34

Ramirez F, Dundar F, Diehl S, Gruning BA, Manke T (2014) deepTools: a flexible platform for exploring deep-sequencing data. Nucleic Acids Res 42: W187–91

Ritter A, Inigo S, Fernandez-Calvo P, Heyndrickx KS, Dhondt S, Shi H, De Milde L, Vanden Bossche R, De Clercq R, Eeckhout D, Ron M, Somers DE, Inze D, Gevaert K, De Jaeger G, Vandepoele K, Pauwels L, Goossens A (2017) The transcriptional repressor complex FRS7-FRS12 regulates flowering time and growth in Arabidopsis. Nat Commun 8: 15235

Robinson MD, McCarthy DJ, Smyth GK (2010) edgeR: a Bioconductor package for differential expression analysis of digital gene expression data. Bioinformatics 26: 139–40

Rosa M, Von Harder M, Cigliano RA, Schlogelhofer P, Scheid OM (2013) The Arabidopsis SWR1 Chromatin-Remodeling Complex Is Important for DNA Repair, Somatic Recombination, and Meiosis. Plant Cell

Rudnizky S, Bavly A, Malik O, Pnueli L, Melamed P, Kaplan A (2016) H2A.Z controls the stability and mobility of nucleosomes to regulate expression of the LH genes. Nat Commun 7: 12958

Rymen B, Sugimoto K (2012) Tuning growth to the environmental demands. Curr Opin Plant Biol 15: 683–90

Sacharowski SP, Gratkowska DM, Sarnowska EA, Kondrak P, Jancewicz I, Porri A, Bucior E, Rolicka AT, Franzen R, Kowalczyk J, Pawlikowska K, Huettel B, Torti S, Schmelzer E, Coupland G, Jerzmanowski A, Koncz C, Sarnowski TJ (2015) SWP73 Subunits of Arabidopsis SWI/SNF Chromatin Remodeling Complexes Play Distinct Roles in Leaf and Flower Development. Plant Cell 27: 1889–906

Salmon-Divon M, Dvinge H, Tammoja K, Bertone P (2010) PeakAnalyzer: genome-wide annotation of chromatin binding and modification loci. BMC Bioinformatics 11: 415

Santisteban MS, Kalashnikova T, Smith MM (2000) Histone H2A.Z regulats transcription and is partially redundant with nucleosome remodeling complexes. Cell 103: 411–22

Schnitzler GR, Cheung CL, Hafner JH, Saurin AJ, Kingston RE, Lieber CM (2001) Direct imaging of human SWI/SNF-remodeled mono- and polynucleosomes by atomic force microscopy employing carbon nanotube tips. Mol Cell Biol 21: 8504–11

Sijacic P, Bajic M, McKinney EC, Meagher RB, Deal RB (2017) Chromatin accessibility changes between Arabidopsis stem cells and mesophyll cells illuminate cell type-specific transcription factor networks. bioRxiv

Sura W, Kabza M, Karlowski WM, Bieluszewski T, Kus-Slowinska M, Paweloszek L, Sadowski J, Ziolkowski PA (2017) Dual role of the histone variant H2A.Z in transcriptional regulation of stress-response genes. Plant Cell

Tabas-Madrid D, Nogales-Cadenas R, Pascual-Montano A (2012) GeneCodis3: a non-redundant and modular enrichment analysis tool for functional genomics. Nucleic Acids Res 40: W478–83

Tian T, Liu Y, Yan H, You Q, Yi X, Du Z, Xu W, Su Z (2017) agriGO v2.0: a GO analysis toolkit for the agricultural community, 2017 update. Nucleic Acids Res

Trapnell C, Roberts A, Goff L, Pertea G, Kim D, Kelley DR, Pimentel H, Salzberg SL, Rinn JL, Pachter L (2012) Differential gene and transcript expression analysis of RNA-seq experiments with TopHat and Cufflinks. Nat Protoc 7: 562–78

Urano K, Kurihara Y, Seki M, Shinozaki K (2010) 'Omics' analyses of regulatory networks in plant abiotic stress responses. Curr Opin Plant Biol 13: 132–8

Valdes-Mora F, Song JZ, Statham AL, Strbenac D, Robinson MD, Nair SS, Patterson KI, Tremethick DJ, Stirzaker C, Clark SJ (2012) Acetylation of H2A.Z is a key epigenetic modification associated with gene deregulation and epigenetic remodeling in cancer. Genome Res 22: 307–21

van Zanten M, Tessadori F, McLoughlin F, Smith R, Millenaar FF, van Driel R, Voesenek LA, Peeters AJ, Fransz P (2010) Photoreceptors CRYTOCHROME2 and phytochrome B control chromatin compaction in Arabidopsis. Plant Physiol 154: 1686–96

Vercruyssen L, Verkest A, Gonzalez N, Heyndrickx KS, Eeckhout D, Han SK, Jegu T, Archacki R, Van Leene J, Andriankaja M, De Bodt S, Abeel T, Coppens F, Dhondt S, De Milde L, Vermeersch M, Maleux K, Gevaert K, Jerzmanowski A, Benhamed M, Wagner D, Vandepoele K, De Jaeger G, Inze D (2014) ANGUSTIFOLIA3 Binds to SWI/SnF Chromatin Remodeling Complexes to Regulate Transcription during Arabidopsis Leaf Development. Plant Cell

Vergara Z, Gutierrez C (2017) Emerging roles of chromatin in the maintenance of genome organization and function in plants. Genome Biol 18: 96

Weber CM, Ramachandran S, Henikoff S (2014) Nucleosomes are context-specific, H2A.Z-modulated barriers to RNA polymerase. Mol Cell 53: 819–30

Weirauch MT, Yang A, Albu M, Cote AG, Montenegro-Montero A, Drewe P, Najafabadi HS, Lambert SA, Mann I, Cook K, Zheng H, Goity A, van Bakel H, Lozano JC, Galli M, Lewsey MG, Huang E, Mukherjee T, Chen X, Reece-Hoyes JS, Govindarajan S, Shaulsky G, Walhout AJM, Bouget FY, Ratsch G, Larrondo LF, Ecker JR, Hughes TR (2014) Determination and inference of eukaryotic transcription factor sequence specificity. Cell 158: 1431–1443

Wigge PA (2013) Ambient temperature signalling in plants. Curr Opin Plant Biol 16: 661–6

Wu JI (2012) Diverse functions of ATP-dependent chromatin remodeling complexes in development and cancer. Acta Biochim Biophys Sin (Shanghai) 44: 54–69

Wu MF, Sang Y, Bezhani S, Yamaguchi N, Han SK, Li Z, Su Y, Slewinski TL, Wagner D (2012) SWI2/SNF2 chromatin remodeling ATPases overcome polycomb repression and control floral organ identity with the LEAFY and SEPALLATA3 transcription factors. Proc Natl Acad Sci U S A 109: 3576–81

Wu MF, Yamaguchi N, Xiao J, Bargmann B, Estelle M, Sang Y, Wagner D (2015) Auxin-regulated chromatin switch directs acquisition of flower primordium founder fate. Elife 4: e09269

Yang S, Li C, Zhao L, Gao S, Lu J, Zhao M, Chen CY, Liu X, Luo M, Cui Y, Yang C, Wu K (2015) The Arabidopsis SWI2/SNF2 Chromatin Remodeling ATPase BRAHMA Targets Directly to PINs and Is Required for Root Stem Cell Niche Maintenance. Plant Cell 27: 1670–80

Yoshida T, Shimada K, Oma Y, Kalck V, Akimura K, Taddei A, Iwahashi H, Kugou K, Ohta K, Gasser SM, Harata M (2010) Actin-related protein Arp6 influences H2A.Z-dependent and -independent gene expression and links ribosomal protein genes to nuclear pores. PLoS Genet 6: e1000910

Zhang D, Li Y, Zhang X, Zha P, Lin R (2016) The SWI2/SNF2 Chromatin-Remodeling ATPase BRAHMA Regulates Chlorophyll Biosynthesis in Arabidopsis. Mol Plant

Zhang D, Li Y, Zhang X, Zha P, Lin R (2017a) The SWI2/SNF2 Chromatin-Remodeling ATPase BRAHMA Regulates Chlorophyll Biosynthesis in Arabidopsis. Mol Plant 10: 155–167

Zhang T, Zhang W, Jiang J (2015) Genome-Wide Nucleosome Occupancy and Positioning and Their Impact on Gene Expression and Evolution in Plants. Plant Physiol 168: 1406–16

Zhang Y, Ku WL, Liu S, Cui K, Jin W, Tang Q, Lu W, Ni B, Zhao K (2017b) Genome-wide identification of histone H2A and histone variant H2A.Z-interacting proteins by bPPI-seq. Cell Res 27: 1258–1274

Zhao M, Yang S, Chen CY, Li C, Shan W, Lu W, Cui Y, Liu X, Wu K (2015) Arabidopsis BREVIPEDICELLUS interacts with the SWI2/SNF2 chromatin remodeling ATPase BRAHMA to regulate KNAT2 and KNAT6 expression in control of inflorescence architecture. PLoS Genet 11: e1005125

